# The Salmon Microbial Genome Atlas enables novel insights into bacteria-host interactions via functional mapping

**DOI:** 10.1101/2023.12.10.570985

**Authors:** Arturo Vera-Ponce de León, Matthias Hoetzinger, Tim Hensen, Shashank Gupta, Bronson Weston, Sander M. Johnsen, Jacob A. Rasmussen, Cecilie Grønlund Clausen, Louisa Pless, Ana Raquel Andrade Veríssimo, Knut Rudi, Lars Snipen, Christian René Karlsen, Morten T. Limborg, Stefan Bertilsson, Ines Thiele, Torgeir R. Hvidsten, Simen R. Sandve, Phillip B. Pope, Sabina Leanti La Rosa

## Abstract

The essential role of the gut microbiota for host health and nutrition is well established for many terrestrial animals, while its importance for fish and particularly Atlantic salmon is unclear. Here, we present the Salmon Microbial Genome Atlas (SMGA) originating from wild and farmed fish both in freshwater and seawater, and consisting of 211 high-quality bacterial genomes, recovered by cultivation (n=131) and gut metagenomics (n=80). Bacterial genomes were taxonomically assigned into 14 different orders, including 28 distinctive genera and 31 potentially novel species. Benchmarking the SMGA, we functionally characterized key populations in the salmon gut that were detected *in vivo*. This included the ability to degrade diet-derived fibers and release vitamins and other exo-metabolites with known beneficial effects, which were validated by *in vitro* cultivation and untargeted metabolomics. Together, the SMGA enables high resolution functional insight into salmon gut microbiota with relevance for salmon nutrition and health.

## Introduction

Efficient and environmentally sustainable aquaculture production systems are urgently required to ensure long-term food security, especially as global seafood consumption is projected to double by 2050 (www.fao.org). For salmonoids, such as Atlantic salmon (*Salmo salar*), this necessitates new ecological sustainable feed ingredients and improvements of broodstock with respect to animal health, feed conversion, and growth. However, an additional layer of complexity that critically influences the path from “feed-to-animal” is the gastrointestinal tract (‘gut’) microbiome. In humans and other vertebrate systems, the gut microbiome has been shown to play a central role in both health and nutrition of its host^1,2^. Decades of research has demonstrated that dietary composition affects the gut microbiome in aquaculture production settings, including in salmon (reviewed in^3,4^). Additionally, since salmon is anadromous, the structure and presumed function of its microbiome is also strongly modulated by whether the fish lives in freshwater (as juveniles) or in seawater (as adults)^5–7^.

To understand the importance of feed-microbiome-host interconnections in salmon and potentially take advantage of these couplings in fish farming, fundamental knowledge gaps must be addressed: namely, how individual microorganisms function, utilize the feed, and interact with each other or the hosts with regards to metabolism and physiology. To date, studies on the gut microbiota in salmon have been based on taxonomic composition of microbial communities via 16S rRNA gene surveys. Accordingly, there is little (if any) genomic sequence information that enable coupling of such compositional data to potential metabolic function or other functional traits in salmon gut microbiomes. Efforts to recover microbial genomes for the salmon gut microbiota have so far been limited to 20 metagenome-assembled genomes (MAGs) that are representatives of dominant *Mycoplasma* populations that constitute a major fraction of the total gut microbiome in adult fish at sea^8,9^. While certain salmon gut samples have indicated *Mycoplasma* spp. levels to be as high as 90%, broad 16S rRNA gene surveys portray much wider diversity that include (and are not limited to) *Aliivibrio*, *Vibrio, Lactobacillus, Photobacterium*, *Carnobacterium, Flavobacterium, Pseudomonas and Psychrobacter* species^7,10–12^. Some of these bacteria have also been recovered using cultivation-dependant approaches^13^, although there has so far been no comprehensive whole genome sequencing study of cultured bacteria from the salmon gut.

The limited success to recover a wide diversity of MAGs from salmon gut samples is likely related to the very low microbial biomass in the fish gut (∼10^4^-10^5^ cells per ml), resulting in host-to-microbiome DNA ratios that typically exceed 9:1^8^. Notwithstanding those alternative approaches, such as single cell genome sequencing and cultivation combined with long read sequencing, could complement traditional shotgun metagenome approaches and result in a more comprehensive database of high-quality and near-complete microbial genomes. In this study, we therefore combine multiple approaches and present the Salmon gut Microbial Genome Atlas (SMGA), a collection of 211 annotated bacterial genomes obtained from the salmon gut microbiota. The SMGA contains genomes from gut microbiota sampled in fish at different developmental stages, in freshwater and seawater, and across farmed and wild populations. We show that the taxonomic profile of these genomes aligns with commonly reported genera that have previously been detected in public 16S rRNA gene surveys of the salmon gut (**Fig. 1**). Lastly, we benchmark and validate the SMGA as a valuable genome reference resource for salmon gut microbiome studies by firstly interpolating putative metabolic functions of keystone populations within an *in vivo* fish trial and then by coupling genomic predictions to culture-based metabolomic analyses (**Fig. 1**).

**Fig. 1.**
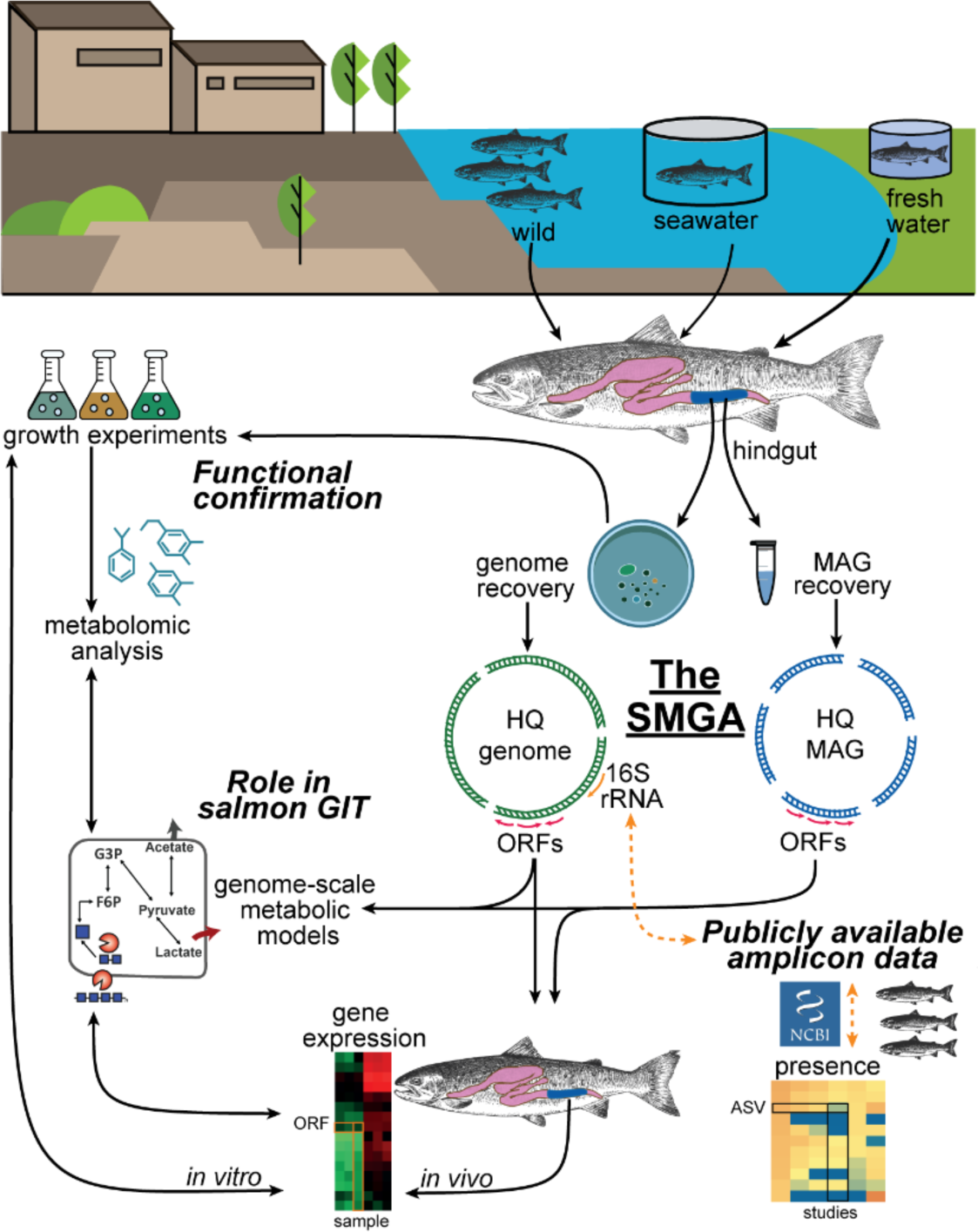
Conceptual overview of the SMGA’s construction and its relevance in connecting the functional potential of the salmon gut microbiota to confirmed metabolic activities. A high-quality (HQ) genome catalogue of the salmon gut microbiota was produced by combining HQ genomes from salmon gut bacterial isolates and HQ MAGs obtained through metagenomics and sorted-cells sequencing. All genomes and MAGs derived from digesta samples were collected from wild fish in seawater as well as farmed fish either from land-based freshwater aquaculture systems or directly from seawater cages. Presence of full-length 16S rRNA gene sequences in the SMGA facilitated the detection of closely related bacteria in publicly available salmon gut-derived amplicon datasets. Open reading frames provided information on the potential metabolic functions and facilitated the mapping of metatranscriptomic (metaT) data derived from salmon feeding trials. Production and consumption of bacterial metabolites presumed from genome-scale metabolic models and from metaT-based reconstruction of active metabolic pathways in key salmon gut bacteria were experimentally validated with *in vitro* growth experiments using the corresponding cultured isolates. This led to confirmation of the beneficial role of the gut microbiota in salmon and uncovers bacterial targets that may be exploited to promote fish physiology and health through dietary interventions.

## Results

### The salmon microbial genome atlas (SMGA): a resource of cultured and uncultured bacteria present in the salmon gut

The recent resurgence of culture-based methods in microbiology has empowered the generation of microbial genome collections that provide valuable connections between phenotype and genotype of microorganisms in a variety of different environments^14–18^. Here, we used different selective media to first culture 71 isolates derived from the midgut of salmons farmed in seawater followed by an additional set of 41 isolates from fish raised in freshwater.

By using Oxford Nanopore long-read sequencing, the genome sequences of the 112 isolates were retrieved and reconstructed as circular chromosomes, and in some cases additional plasmids were recovered in these genomes (**Supplementary Table S1**). Together, these 112 sequenced isolates from the midgut of Norwegian Atlantic salmon constitute the Norwegian Atlantic Salmon Gut Bacteria Culture Collection (NAS-GBCC; **Supplementary Table S1**). The isolates were cryopreserved and are currently available upon request from the Norwegian University of Life Sciences.

Taxonomic classification using GTDB-Tk showed that the majority (n=91) of these genomes affiliated to the Pseudomonadota (Proteobacteria) phylum, followed by the Bacillota (Firmicutes) and Bacteroidota phyla (n=10 and n=9). Using an operational species definition based on genome similarity, i.e. a 95% average nucleotide identity (ANI) threshold^19,20^, genome phylogeny and GTDB-Tk analysis^20^ showed that 35 isolates represent putative novel species among the genera: *Aliivibrio*, *Flavobacterium, Glutamicibacter, Photobacterium, Pseudomonas, Psychrobacter, and Shewanella* (**Fig. 2** and **Supplementary Table S1**). Species novelty was additionally supported with established 16S rRNA gene sequence identity of 98.7-94.5% using available genomes in NCBI^21^. We further supplemented the resulting genome collection with 19 previously published genomes of gut-derived *Latilactobacillus* isolates from salmons farmed in Norway and North America^22^, amounting to a total of 131 genomes of isolated strains.

**Fig. 2.**
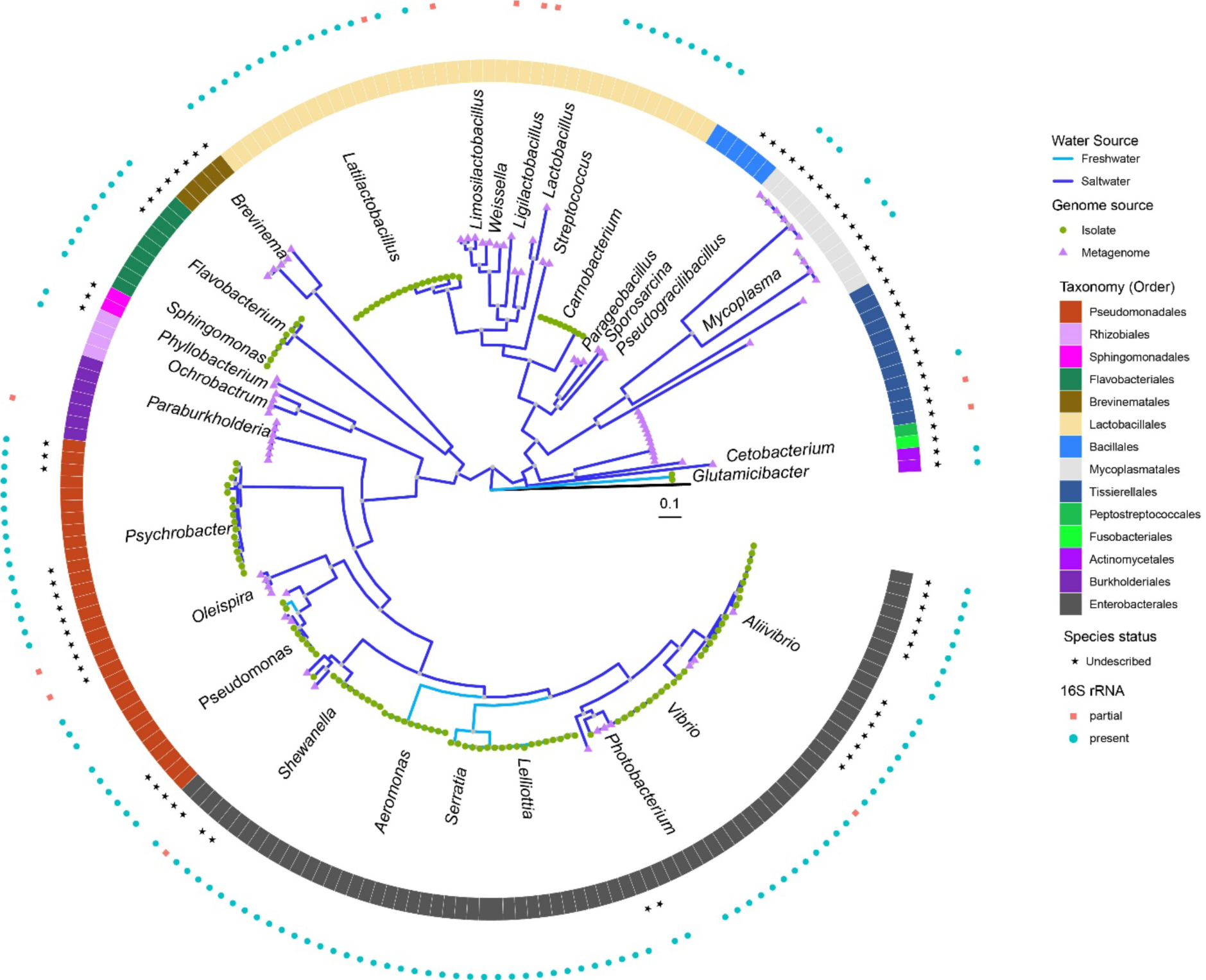
Phylogenetic tree showing the diversity and source, of the recovered bacterial genomes and MAGs. The cladogram depicts the taxonomic classification of all the 211 SMGA genomes coloured by order (inner ring). Grey dots in the cladogram indicates a Bootstrap support higher than 70 %. A green dot represents a genome from a cultured isolate while a purple triangle indicates a MAG. Genomes associated to undescribed species are indicated with a star (middle ring), while genomes encoding a partial or complete 16S rRNA gene operon are indicated by red squares and light blue circles, respectively (outer ring). Sample source is depicted with either a light blue or a dark blue branch for freshwater or seawater salmon, respectively. The genome of *Prochlorococcus marinus* subsp*. marinus* str. CCMP1375 (RefSeq GCF_000007925.1) was used as an outgroup (black branch). Scale bar indicates 10% estimated sequence divergence.

To broaden the diversity of our genome collection, we incorporated as-yet uncultured microorganisms by producing approximately 1.2 Tbp of shotgun data from 93 samples derived from the gut content of salmons farmed both in fresh- and seawater. Notably, mapping of our raw metagenomic reads to the salmon genome revealed a 90.5-99.2% fraction originating from the host, which was higher than in previous studies^8^. In response to high levels of host DNA contamination we also sequenced metagenomic DNA isolated after host DNA depletion, which resulted in a lower fraction of reads mapped to the host (24.2-71.9%). Furthermore, fluorescence-activated cell sorting (FACS) was applied to partition microbial cells from debris and host cells and subsequently sequence pools of such cells as “mini-metagenomes”. Assembly of the different metagenomes after removal of host reads, followed by binning of the assembly output, resulted in 68 MAGs fulfilling quality criteria for medium to high-quality MAGs (estimated >50% completeness and <5% contamination, according to the standards described in^23^). Only three of these MAGs were obtained from mini-metagenomes (**Supplementary Table S1**), reflecting the difficulties to separate out microbial cells from complex, low microbial biomass samples. We additionally assembled two MAGs from previously published metagenomes obtained from gut samples of salmons also farmed in Norway^24^. Finally, we included 10 previously published medium to high-quality MAGs, which had been derived from gut samples of wild salmons caught along the coast of northern Norway^9^. Taken together, the SMGA thereby feature a collection of 80 MAGs, including 31 of high-quality (>90% completeness, <5% contamination) according to the standards described in the “Minimum Information about a Metagenome-Assembled Genome” (MIMAG)^23^. The MAGs significantly increased the taxonomic diversity of the SMGA. Besides Lactobacillales (14 MAGs), Enterobacteriales (9 MAGs) and Pseudomonadales (7 MAGs) which were frequent also among the isolates, nine orders were solely represented by MAGs, with Mycoplasmatales (12 MAGs), Tissierellales (12 MAGs), Burkholderiales (7 MAGs) and Bacillales (6 MAGs) being most frequent.

In total the SMGA consists of 211 genomes and MAGs (**Supplementary Fig. S1**), including 31 undescribed species, and comprises a total of 286,891 unique protein-coding genes (and 739,323 non unique protein-coding genes). At 95% ANI (average nucleotide identity) threshold, genomes and MAGs grouped into 62 species-like clusters (mOTUs), with pan-genomes comprising up to 27,640 unique proteins (**Supplementary Fig. S2**). In general, genomes grouped distinctly based on whether they were isolated from freshwaters (e.g. *Lelliottia* and *Serratia* spp.) or marine systems (e.g. *Photobacterium* and *Mycoplasma* spp.). There were nevertheless taxa that were observed in both, such as *Carnobacterium* and *Pseudomonas* (**Fig. 2**).

### Benchmarking the value of SMGA - linking 16S data to complete microbial genomes

Our combined use of short and long-read DNA sequencing ensured that 129 genomes and 17 MAGs from the SMGA encoded full-length 16S rRNA genes, which enabled searches for the occurrence of SMGA bacteria in the plethora of amplicon datasets that dominate the salmon gut microbiome literature. At a 97% identity cut-off when comparing SMGA 16S rRNA sequences to amplicon sequence variants (ASVs), 144 out of 146 SMGA bacteria were detected in publicly available 16S rRNA gene datasets from either *in vivo* trials or *in vitro* models with salmon gut microbial communities, as well as datasets generated within this study (ImpTrial1 and ImpTrial2) (**Supplementary Fig. S3**)^10,25–34^. SMGA bacteria were detected in microbiomes not only from Norwegian salmon populations, but also in gut samples from wild and farmed Atlantic salmon retrieved from Scotland^27^, the UK^28^ and Chile^34^. Our findings validate that 99% of the SMGA microbes that featured complete 16S rRNA genes are also found in a wider range of salmon gut microbiomes. This included prevalent genera such as *Carnobacterium*, *Lactobacillus*, *Flavobacteria*, *Photobacterium*, *Shewanella*, *Vibrio* and *Aliivibrio* that are routinely observed in salmon microbiome research^7,10–12^. Using the reverse approach, 16S rRNA gene amplicon data from taxonomic surveys can be linked to genome encoded functional traits from the SMGA’s collection to provide additional metabolic and functional context. This will eventually also enable cross-study comparisons and aid the prediction of microbiota functions potentially resulting in growth-related and health-related metabolites beneficial for the salmon host.

### Putative metabolic capabilities encoded in the SMGA bacterial genomes

Equipped with our genome inventory, we subsequently explored the metabolic potential of the individual strains using functional annotation databases (**Fig. 3a**). We also generated genome-scale models for 94 of the 211 genomes (Methods, **Supplementary Fig. S4-S5, Supplementary Table S2**), three of which were validated against exo-metabolomic data (Methods, **Supplementary Fig. S6-S7, Supplementary Table S3).** We used these metabolic models to predict metabolic fluxes and metabolite exchange, which we also used for the exploration of the metabolic potentials. As expected, core metabolic pathways (glycolysis, etc), glucose consumption and acetate metabolism were largely similar among strains. Both facultative and strict aerobes were identified, and fittingly respiration and fermentation were predicted across the SMGA genomes. More specifically, some strains presented genes and pathways that could lead to potential beneficial metabolites in the salmon gut such as short chain fatty acids, amino acids as well as B- and K-vitamins (**Supplementary Fig. S4**). For example, lactic and succinic acid production was predicted via genome annotation for many Pseudomonadota and Bacillota and was further supported by prediction of lactate and succinate metabolite exchange (**Supplementary Fig. S5**). Nitrogen cycling varied across the SMGA genomes, with various Pseudomonadota species predicted to either take up nitrate, excrete nitrite or perform dissimilatory nitrate reduction to ammonium (e.g. *Allivibrio* and *Shewanella* spp.). Metabolism of ammonium, ornithine and citrulline via the urea cycle was also predicted for certain bacteria, including *Pseudomonas* and *Carnobacterium* spp. Metabolism of amino acids such as glycine, alanine, leucine, valine, aspartate and arginine varied considerably in their predicted uptake and excrement, highlighting metabolic points of difference across the SMGA.

**Fig. 3.**
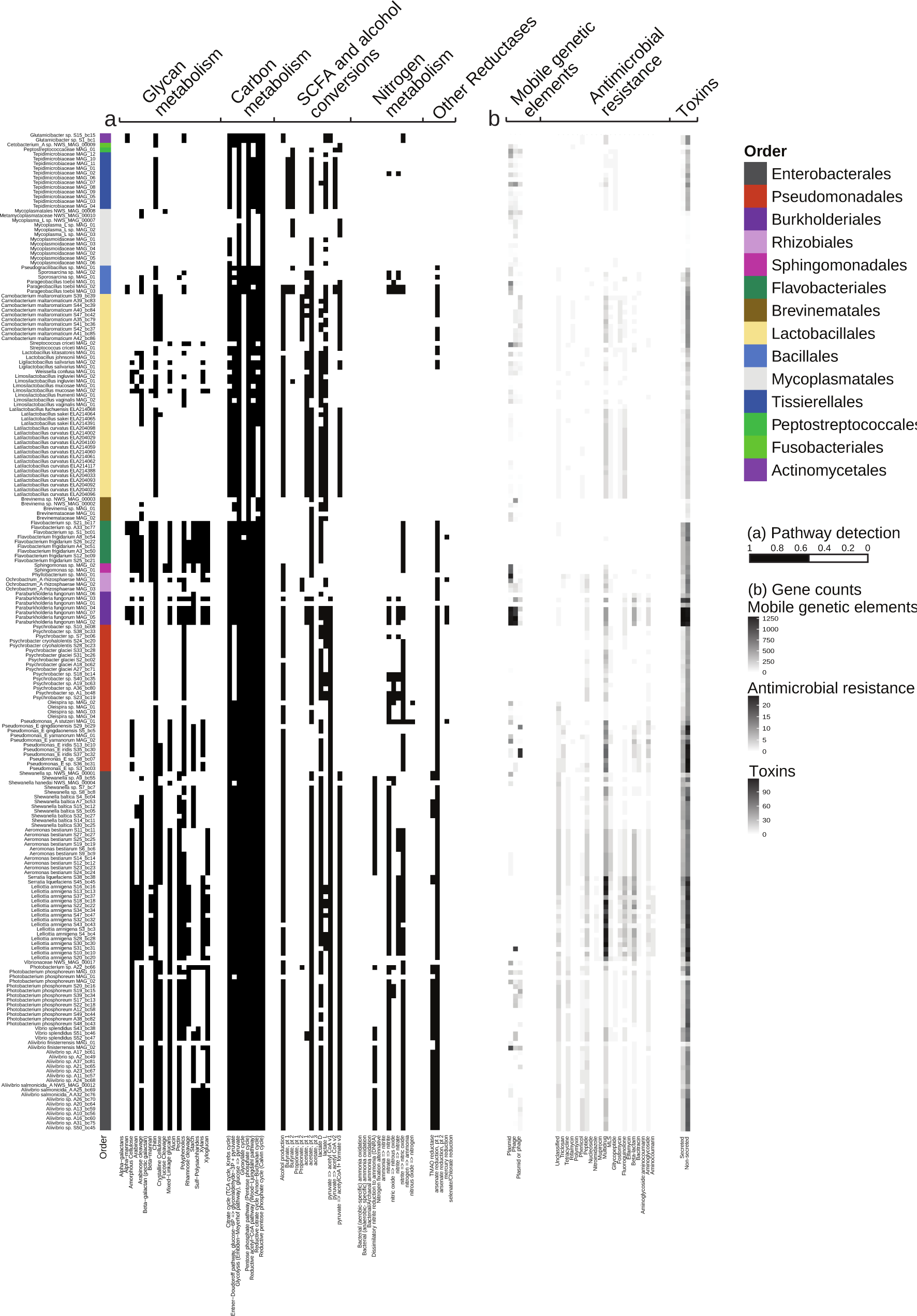
Metabolic functions encoded by the 211 genomes in the SMGA. **a.** Heatmap showing the presence of genes/pathways (listed on the lower x-axis) across various functional categories (listed on upper x-axis) found in each genome (y-axis). The presence of a gene/pathway is denoted by a black box and considered present if >50% of the genes in the DRAM module are encoded. Genes/pathways that are not detected are represented by a white box. DRAM functional categories, sub-categories and functional IDs are listed in **Supplementary Table S4**. **b**. Number of genes putatively encoding potential pathogenic factors harbored by the SMGA, including factors for adherence, motility, toxins and antimicrobial resistance.

Aquaculture practices are continuously evolving and with that also shifts in preferred salmon diets. Such management practices can benefit from insights about the ability of salmon gut microbiota to access and metabolize marine-, terrestrial plant- and insect-derived carbohydrates. In this context, the SMGA was also assessed for the presence of carbohydrate active enzymes (CAZymes) such as glycoside hydrolases (GH) and polysaccharide lyases (PL) (**Fig. 3a, Supplementary Fig. S8** and **Supplementary Table S4**). Most of the genomes encoded CAZymes, with an average number of 115 genes per genome. MAGs with the smallest number of CAZymes were affiliated with the order Mycoplasmatales (average number of CAZyme = 7, **Supplementary Table S4**), which is in line with previous studies^8,9^, and imply their limited contribution to metabolism of dietary components in salmon. The dominant CAZy family within the SMGA was GH13, which has been proven to have the capacity to metabolize starch^35^. Notably, prevalent enzymes in Enterobacterales, Lactobacillales and Pseudomonadales were GH18 and GH19 (followed by GH20), which enable microorganisms to depolymerize chitin through the hydrolytic utilization pathway^35^. Intriguingly, AA10 LPMOs, that have been shown to be involved in an alternative (oxidative) chitin utilization pathway^36^, were detected in the genomes of Enterobacterales, Lactobacillales and a few Pseudomonadales. CAZymes involved in utilization of terrestrial and marine plant-derived carbohydrates (e.g. beta-mannan, beta-glucans, xylans, cello-oligosaccharides, manno-oligosaccharides and algal polysaccharides) included GH1, GH2, GH3, GH5, GH8, GH9, GH10, GH16, GH26, GH36, GH43 and GH94 among others^35^. In addition, CAZymes belonging to the families GH28, GH35, GH78, GH105, GH147, PL2, PL9 and PL22 for deconstruction of the plant pectic polysaccharide rhamnogalacturonan-I were detected in some Enterobacterales and Lactobacillales genomes^35^. A few Pseudomonadota and Bacteroidota genomes harboured genes encoding CAZymes for depolymerization of host mucin-derived oligosaccharides, including GH29, GH33, GH109, GH112 and GH129^35^.

Virulence arising from the salmon gut microbiome can impact fish health. Hence, we screened the genomes for elements encoding virulence factors, bacterial toxins, and antimicrobial resistance in the 211 genomes and MAGs of the SMGA (**Fig. 3b**). Mobile genetic elements, including both phage- and plasmid-derived sequences, were detected in 147 SMGA genomes/MAGs. Genes encoding putative toxins were identified in all SMGA genomes and MAGs using PathoFact searches^37^ against the Toxin and Toxin Target Database (T3DB)^38,39^. Of these putative toxins, 27% and 73% were predicted to be secreted and non-secreted pathogenic factors, respectively. The highest numbers of toxin genes were found in Pseudomonadota and particularly the genera *Paraburkholderia*, *Pseudomonas*, *Serratia* and *Lelliottia*, with up to 233 such genes in one *Paraburkholderia* MAG, of which 51% were predicted to be secreted. With regards to gene categories enabling antimicrobial resistance (AMR), the majority of the predicted resistance genes included beta-lactam-, fluoroquinolone-, aminoglycoside-, tetracycline-, peptide- and multidrug-resistance. Genes associated with beta-lactam resistance were found to be particularly enriched in the genera *Lelliottia*, *Serratia*, *Pseudomonas* and *Aeromonas*, suggesting they might include potentially pathogenic members.

### The SMGA facilitates functional understanding of the salmon gut microbiome

To showcase how the SMGA can be used to overcome one of the most obtrusive knowledge gaps in salmon gut microbiome research, namely the lack of functional understanding, we used the SMGA as a database to map metatranscriptomes from gut samples obtained from growing fish fed a standard commercial diet and collected at different life stages. This integrated omic approach facilitated detection of 116,888 expressed genes (15.81% Identification Rate) and, for the first time, identified several functionally active microbial populations *in vivo*. Expressed metabolic pathways varied among the 4 life stages sampled, with the majority of expressed genes (**Supplementary Fig. S9a, Supplementary Table S5**) mapping to Enterobacterales (n=36,026), Pseudomonales (n=32,649), Lactobacillales (n=18,177), Burkholderiales (n=13,292) and Tissierellales (n=8,977). Bacteria belonging to these taxa are commonly encountered in 16S rRNA gene amplicon surveys, suggesting that they likely play active roles in the salmon gut microbiome. Lactobacillales were one of the most active populations at all the life stages sampled, with a large proportion of expressed genes in gut samples collected from salmon in seawater. Based on the expressed CAZymes, Lactobacillales metabolized starch and maltose (through GH13s, GH65s and GH126s), mannose- and cello-oligosaccharides (through GH1s, GH2s, GH3s), and potentially polymeric β-mannan and cellulose (through GH5s) (**Fig. 4a**, **Supplementary Table S6**).

**Fig. 4.**
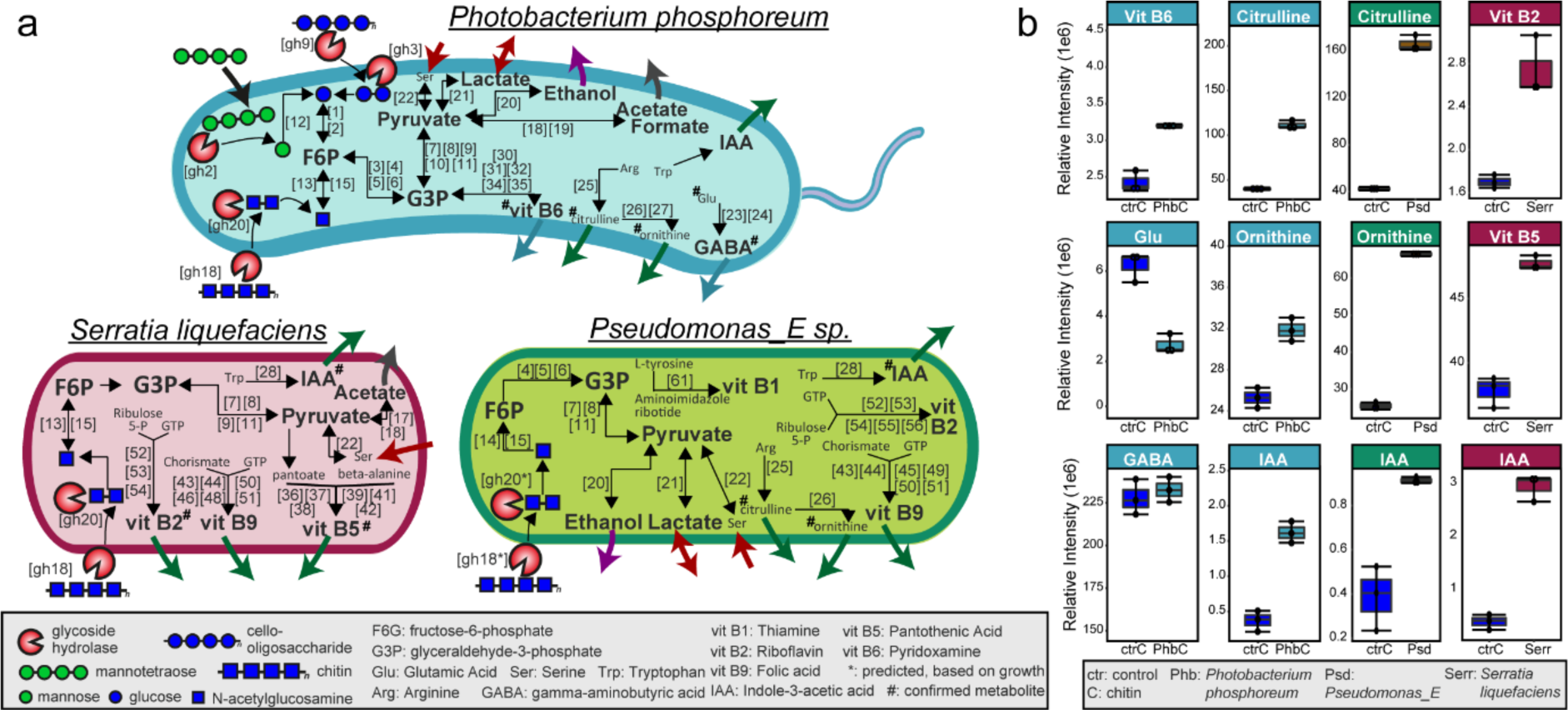
Schematic overview of active metabolic pathways detected in representative species of the SMGA based on metatranscriptomics, *in vitro* growth experiments and untargeted metabolomic of spent supernatants. **a.** Cartoons depicting detected active metabolic pathways of *Photobacterium phosphoreum*, *Serratia liquefaciens*, and *Pseudomonas_E* using metatranscriptomics data generated from hindgut samples obtained from fish farmed in fresh and seawater under a standard commercial diet. Genes are displayed as boxed numbers and are fully listed in **Supplementary Table S5**. Uptake and release of metabolites, amino acids and dietary fibers is indicated with arrows. **b.** Untargeted metabolomics of supernatant samples after growth of *P. phosphoreum*, *S. liquefaciens*, and *Pseudomonas* in a medium supplemented with chitin. Bars represent median and boxes interquartile range determined from 3 independent replicates. Values for control and experimental samples are in **Supplementary Table S7**.

Given the opportunity to validate omic-based inferences with cultivation experiments, we further explored the gene expression profile of three bacteria present in our culture collection. *Photobacterium phosphoreum* S39_bc34 was identified as the bacterium expressing the highest number of genes during the seawater stage (Supplementary Fig. S9b). This included genes encoding CAZymes involved in depolymerization of chitin (GH18, GH20), hemicelluloses (manno- and cello-oligosaccharides, including GH2, GH3, GH9 and GH92; arabinogalactan, including GH154 and PL22), starch (GH13) and glycosaminoglycans (PL8) (**Fig. 4a**, **Supplementary Table S6**), suggesting that this bacterium is capable of readily utilizing these abundant biopolymer substrates. In all salmon gut contents, *P. phosphoreum* S39_bc34 showed high expression of genes coding for enzymes inferred in energy generation through glycolysis and pyruvate metabolism, with potential formation of acetate and formate. These observations were consistent with the metabolic model-based predictions (**Supplementary Fig. S4-7**, **Supplementary Table S3**). Based on the gene expression data, we also predicted several pathways involved in amino acid metabolism, with an active L-serine dehydratase [EC:4.3.1.17; VMH ID: r0060] for the conversion of serine into pyruvate; active arginine deiminase [EC:3.5.3.6; VMH ID: ARGDA], ornithine carbamoyltransferase [EC:2.1.3.3; VMH ID: OCBT] and carbamate kinase [EC:2.7.2.2; VMH ID: CBMKr] which have demonstrated activity on arginine conversion into citrulline and ornithine that can be further exchanged with putrescine using an active putrescine:ornithine antiporter^40,41^. Accordingly, the metabolic model could secrete ornithine and putrescine (**Supplementary Table S3)**. Expression of genes inferred in glutaminase [EC:3.5.1.2; VMH ID: GLUN] and glutamate decarboxylase [EC:4.1.1.15; VMH ID: GLUDC] activity for conversion of glutamic acid to gamma aminobutyric acid (GABA), followed by extracellular export of GABA through an active glutamate:GABA antiporter^42^, were also detected (**Fig. 4a**, **Supplementary Table S6**). Expressed genes encoding enzymes involved in the route that joins glyceraldehyde 3-phosphate and D-ribulose 5-phosphate for *de novo* synthesis of pyridoxamine (vitamin B6) were also found^43^, presumably resulting in the release of this vitamin in the salmon gut (**Fig. 4a**, **Supplementary Table S6**).

Focusing on bacteria from our culture collection that were highly active in the gut of salmon during the freshwater phase, *Serratia liquefaciens* S38_bc38 and *Pseudomonas_E* sp. S3_bc03 expressed genes for hydrolysis of chitin and chito-oligosaccharides (GH18 and GH20), manno- and cello-oligosaccharides (GH2, GH3,), starch (GH13, GH15 and GH31) and pectic oligosaccharides (PL22), all resulting in the release of monosaccharides that can subsequently be metabolized through an active glycolytic pathway (**Fig. 4a**, **Supplementary Table S6**). Expressed genes encoding enzymes involved in metabolism of serine (arginine deiminase [EC:3.5.3.6; VMH ID: ARGDA]), arginine (arginine deiminase [EC:3.5.3.6], ornithine carbamoyltransferase [EC:2.1.3.3; VMH ID: OCBT] and carbamate kinase [EC:2.7.2.2; VMH ID: CBMKr]) and tryptophan metabolism (acetaldehyde dehydrogenase [EC:1.2.1.10; ACALD]) were detected in *S. liquefaciens* S38_bc38 and *Pseudomonas_E* sp. S03_bc03. Some but not all the reactions associated with these genes were also present in the respective metabolic models (**Supplementary Table S3**). Expressed pathways for vitamin metabolism were detected in *S. liquefaciens* S38_bc38, presumably producing pantothenate (vitamin B5) from 2-dihydropantoate and b-alanine, riboflavin (vitamin B2) from guanosine triphosphate (GTP) and D-ribulose-5-phosphate as well as folate (vitamin B9) from GTP and chorismate (**Fig. 4a**, **Supplementary Table S6**). In *Pseudomonas_E* sp. S03_bc03, besides active genes for production of vitamin B2 and B9, we detected expressed genes that are part of the thiamine (vitamin B1) biosynthetic pathway^43^. Overall, these results demonstrate the power of applying the SMGA as a genome database to map realized functions at the level of individual strains and allowed us to show that salmon gut microbes can supply the host with beneficial metabolites, including short chain fatty acids, vitamins and polyamines.

To validate our omic-based functional inferences from fish trials, we further functionally characterized a selection of our cultured isolates. The isolates *P. phosphoreum* S39_bc34, *S. liquefaciens* S38_bc38 and *Pseudomonas_E* sp. S3_bc03 were chosen and their production and/or consumption of metabolites were compared when each individual microbe was grown on a medium supplemented with chitin against a negative control. Consistent with the *in vivo* RNAseq-based predictions (**Fig. 4b**), all isolates grew on chitin in liquid monocultures. Untargeted metabolomic analyses of the spent culture media from *P. phosphoreum* confirmed the presence of metabolites derived from amino acids catabolism, with decreased levels (consumption) of glutamic acid, arginine and serine and increased levels (production) of GABA, citrulline, ornithine, and pyridoxamine (**Fig. 4b**). Production of indole-3-acetic acid (3-IAA), a metabolite from tryptophan catabolism, was detected (**Fig. 4b**), although this pathway remains uncharacterized in *P. phosphoreum*. As predicted based on the *in vivo* RNAseq data, *P. phosphoreum* produced pyridoxamine (**Fig. 4b**), likely from intermediates of glycolysis and the pentose phosphate pathway, while both *S. liquefaciens* produced B-vitamins, including riboflavin, panthotenic acids and pyridoxamine (**Fig. 4b**). We also found an increase of 3-IAA acid in the spent media of *S. liquefaciens* and *Pseudomonas_E*, compatible with tryptophan catabolism (**Fig. 4b**). Synthesis of citrulline and ornithine from the arginine pathway was confirmed in *Pseudomonas_E* (**Fig. 4b**), and the corresponding metabolic models predicted that all three strains could take up lactic acid and serine, if provided in the medium (**Supplementary Fig. S6, Supplementary Table S3**). Together, these findings demonstrate the value of the SMGA resource by facilitating the identification and experimental validation of active metabolic pathways and metabolites in salmon gut bacterial species.

## Discussion

A critical obstacle for comprehensive functional understanding of the salmon gut microbiome for physiology and nutrition, is the current lack of a genome-centric view which captures the diversity and metabolic potential of the resident microbial populations. This knowledge is pivotal for identifying active metabolism of gut microbes and for translating alterations in the abundance of gut microbial populations into metabolic change with the potential to alter gut function. Here we present the hitherto most extensive microbial genome dataset from Atlantic salmon, obtained through cultivation and whole genome sequencing, as well as metagenomic sequencing of samples from gut contents of wild and farmed fish both in fresh and sea water (**Fig. 1**). The atlas comprises 131 genomes and 80 MAGs, and a total of 286,891 unique proteins.

We demonstrate that the catalogue is widely detected in the salmon gut, as 99% of the SMGA genomes with a complete 16S rRNA gene can be found in publicly available salmon gut amplicon sequencing datasets (**Supplementary Fig. S3**). To showcase the strength of this resource for mechanistic studies of the salmon gut microbiome, we functionally annotated all genomes, created predictive genome-scale metabolic reconstructions and mapped metatranscriptomic data from an independent fish trial to the SMGA. We further confirmed our metabolic predictions with actual experimentally measured metabolites from selected pure bacterial cultures. This multi-faceted approach generated insights into several key populations, that are routinely detected in 16S rRNA gene-based microbiome surveys, in the metabolism of dietary components, including chitin, cello-oligosaccharides and manno-oligosaccharides (**Fig. 4**).

Chitin is a common carbohydrate found in the natural diet of Atlantic salmon as a component of the exoskeleton of insects and crustaceans^12^. While Atlantic salmon possess endogenous chitinases that have some level of activity for use and exploitation of this dietary component^44,45^, our findings show that gut Gammaproteobacteria (*P. phosphoreum*, *S. liquefaciens* and *Pseudomonas*) can efficiently degrade chitin in pure culture while *in vivo* they express chitinases and are inferred as active degraders that contribute towards break-down of this polymer (**Fig. 4**). Previous work, based on amplicon sequencing, has shown that dietary inclusion of chitin-rich insect meals modulates the composition of gut microbiota in salmon post-smolt, leading to increased relative abundance of gram-positive Actinomycetota and Bacilli^12^ and decreased abundance of Pseudomonadota. Similarly, a study from Li et al.^10^ reported an enrichment of Bacilli in the gut of salmon fed an insect meal diet, although the authors highlighted that the composition of the gut microbiota closely resembled that of the feed. It may be possible that reports of dietary effects of chitin-rich feeds are in part biased by the fact that the 16S rRNA amplicon sequencing-based approach provides a community overview but does not discriminate between metabolically active and inactive populations in the gut (including carry-over of microbial DNA from abundant microbial populations in the feed); as a consequence of this, the observed changes in microbiome composition may not directly translate into functional alterations that impact host’s metabolism. This challenge can be addressed by using the SMGA as a reference database for metatranscriptomic studies to clearly discriminate metabolic activities as well as to decrypt microbial mechanisms for chitin degradation and their implications on salmon nutrition and physiology.

From an industry perspective plant-based feed ingredients are increasingly being used as an alternative to fish-based meals. In this context, active pathways for the deconstruction of plant-derived fibers, fermentation of the resulting sugars, and production of acetate and formate, were detected in *P. phosphoreum,* a member of the core gut microbiota in both healthy farmed as well as wild Atlantic salmon during seawater stages^46^ (**Fig. 4**). Of further nutritional importance, we observed an upregulation of genes for B-vitamin related enzymes in several active microbial populations. Production of pyridoxamyne (vitamin B6) was indeed detected in *P. phosphoreum* from pure cultures and *in vivo* trials, while *S. liquefaciens* and *Pseudomonas* were found to be involved in both *in vitro* and *in vivo* production of vitamin riboflavin (B2), pantothenic acid (B5) and folic acid (B9) (**Fig. 4a**). Provision of microbially-derived B vitamins has been shown to be important for development and survival of a variety of animal hosts^43,47^. In essence, microbes supply vitamins that are limited in the diet or complement diet provision, ensuring growth in scarce dietary conditions. In Atlantic salmon, B-vitamins such as folate, riboflavin, niacin, pyridoxamine, and cobalamin have been shown to have effects on hepatic transcriptional and epigenetic regulation of pathways related to lipid metabolism^48^, while increased amounts of pyridoxamine have been associated with improved fish health^49^.

Pathways for metabolism of amino acids, including glutamic acid, arginine and tryptophane, were also found to be up-regulated in all the most active bacteria in the salmon gut, with production of GABA, ornithine, citrulline and 3-IAA confirmed via culture experiments and metabolomic analysis (**Fig. 4b**). Ornithine and citrulline have a central role in arginine metabolism and have been previously associated with increased fish growth, although the underlying mechanism is not yet known^50,51^. While the beneficial role of GABA and 3-IAA for host physiology has been documented in humans where the former induce a calming effect and the later improve intestinal mucosal barrier functions^42,52,53^, the physiological mechanism in Atlantic salmon has not been shown. Collectively, our results reveal hitherto unknown aspects of microbial fermentation, amino acid metabolism and vitamin provision and strengthen the knowledge on the involvement of the gut microbiome as a continuous source of beneficial metabolites that support health and growth in salmon.

In conclusion, our findings provide valuable new insights related to carbohydrate, amino acid and vitamin metabolism in the salmon gut microbiota and reveal that gut bacteria can potentially affect host physiology through provision of several beneficial metabolites. Further, our work establishes a valuable genomic resource that can serve as a reference for genome-resolved functional omics to evaluate the metabolic potential and actual activity of key microbial players in the salmon gut under varying experimental conditions. Finally, we exemplify the *in vivo* detection and in-depth *in vitro* characterization of four bacteria and showcase how the SMGA can readily facilitate major conceptual advances regarding microbial metabolic capacities in the salmon gut and empower new research efforts to shed light on microbiome functions, dynamics and metabolic interactions with the salmon host.

## Materials and Methods

### Bacterial isolation and cultivation

For strains isolated at NMBU, samples were obtained from adult Atlantic salmon kept at the Center for Fish Research, NMBU, Ås, Norway. The fish were reared in recirculated freshwater tanks (14.4 ± 0.4 °C) and kept under continuous light. Mid-gut contents were collected from six salmons into sterile 15 mL Eppendorf tubes using aseptic techniques. Samples were immediately homogenised in sterile PBS (1:1 w/v) and a 1:10 dilution series performed. Hundred mL of each dilution was then plated onto BHI (Brain Heart Infusion, Oxoid) agar supplemented with 2.5% NaCl and TSB (Tryptone Soy Broth, Thermo Scientific) agar supplemented with 2.5% NaCl and 5% glucose. Plates were incubated at 15 °C for 3-7 days prior. Pure cultures were obtained by picking individual colonies and re-streaking onto fresh plates. This process was repeated until purity was achieved. Fifty-six of these isolates were taxonomically classified by amplifying the full-length 16S rRNA gene using the primers 27F (5′-AGAGTTTGATCATGGCTCA-3′) and 1492R (5′-TACGGTTACCTTGTTACGACTT-3′), followed by Sanger sequencing (Eurofins Genomics) with both primers. Sequences were analyzed and edited in BioEdit and BLASTed against the sequences available in GenBank. Colonies of interest were cultured on the appropriate liquid culture and genomic DNA was extracted using the Nanobind CBB Big DNA Kit (Circulomics) according to the manufacturer’s guidelines, using the high-molecular weight (HMW) protocol for gram-positive bacteria. The DNA concentration was measured on a Qubit 3.0 fluorometer with the Qubit dsDNA BR assay kit (Thermo Fisher Scientific) and the DNA quality was measured by gel electrophoresis on an BioRad Gel Doc EZ Imager (Bio-Rad Laboratories, Inc.).

For bacteria isolated at NOFIMA (Norway), samples were obtained from adult Atlantic salmon kept in seawater cages. Mid-gut contents were processed as described above, with the exception that appropriate dilutions were plated onto LB (Sigma-Aldrich) agar supplemented with 2.5% NaCl or MacConkey (Merck KGaA, Germany) agar plates. Seventy-six isolates were identified using 16S rRNA gene sequencing as described above. Selected isolates were grown in liquid cultures and HMW genomic DNA for sequencing was obtained using the procedure described above.

### Genome sequencing and assembly of bacterial isolates

Long-fragment DNA sequencing was conducted using an Oxford Nanopore Technologies (ONT) PromethION sequencer. The sequencing libraries (one with DNA from the 56 strains isolated at NMBU and one with DNA from the 76 strains isolated at NOFIMA) were prepared using the 1D Native barcoding kit EXP-NBD196 (ONT), followed by Oxford Nanopore 1D Genomic DNA by ligation sequencing kit SQK-LSK109 (ONT), according to the manufacturer’s instructions. The total eluted library was then loaded onto an ONT FLO-PRO002 R9.4.1 flow cell, following the manufacturer’s guidelines, and sequenced for 48 h on a PromethIon device using the MinKNOW v.4.0.5 software. Basecalling was performed with Guppy v.4.0.11 in “high-accuracy basecalling” mode. After basecalling, reads were filtered by quality using FiltLong v0.2.1 (GitHub - rrwick/Filtlong: quality filtering tool for long reads) with the --min_length 1000 and -- keep_percent 90 parameters. Filtered reads were then assembled using Fly^54^ in “-- nano-raw” mode using default parameters. Quality of the bins was assessed using BUSCO v1.0 with the “-m geno –auto-lineage-prok” parameters^55^.

### Sample collection for metagenomics and Fluorescence-activated cell sorting (FACS)

Approximately 300 mg hindgut content were collected from dissected salmon, dissolved in 1 ml PBS with 150 µl GlyTE buffer in a cryotube and stored at −80°C until processing. Three different methods were used to potentially enrich bacteria from hindgut samples before cell sorting. (I) Filtration using 1.6, 2.7 or 5 µm syringe filters. (II) Extracting the supernatant after centrifugation. Different times (5 sec – 5 min) and centrifugation speeds (500 x g – 5 000 x g) were tested. (III) Extracting different fractions from Nycodenz density gradient centrifugations at 10 000 x g for 40 min. Different Nycodenz concentrations between 45 and 80% were tested. Cell sorting was performed at the SciLifeLab Microbial Single Cell Genomics Facility with a MoFlo™ Astrios EQ sorter (Beckman Coulter, USA). Cells were stained with SYBR Green I Nucleic Acid stain (Invitrogen™, Thermo Fisher Scientific, MA, USA) and sorted based on forward scatter and fluorescence intensity at 488/530 nm excitation/emission into 384-well plates by collecting 1 - 200 events per well using a 70 or 100 µm nozzle. The Phi29 enzyme was used for whole genome amplification via multiple displacement amplification (MDA). SYTO 13 nucleic acid stain was added to the reaction to monitor DNA amplification over time. Wells were screened for bacterial cells by 16S rRNA gene amplification using primers Bakt_341F and Bakt_805R^56^.

### Extraction of DNA for shotgun metagenomics

Crude metagenomes were extracted from hindgut samples using the DNeasy PowerSoil Pro Kit (Qiagen). To obtain host-depleted metagenomes, the HostZERO Microbial DNA Kit (Zymo) was used with the following protocol adjustments. Hindgut samples were centrifuged at 13 000 x g for 5 min, supernatants removed, pellets resuspended in 1.5 ml PBS buffer and transferred to 5 ml tubes containing ∼0.7 ml sterile 2 mm glass beads. To mostly lyse host cells but preserve microbial cells, the tubes were agitated in a FastPrep24 instrument (MP Biomedicals, Santa Ana, CA, USA) for 2 x 30 sec with a 30 sec break in between at speed level 3.4. Tubes were then centrifuged at 16 000 x g for 2 min, supernatant removed, pellet resuspended with 2 ml Host Depletion Solution (Zymo) and transferred without the beads to two 1.5 ml tubes per sample. From here, the HostZERO Microbial DNA Kit protocol was followed, whereby each sample was pooled again into one tube at the step of adding 100 µl Microbial Selection Buffer. Extracted DNA of at least nine samples needed to be pooled to obtain 5-10 ng DNA for library preparation.

### Genome-resolved metagenomes

10 MAGs were obtained from a previous study of gut microbes from wild-salmon populations^9^. A total of 70 MAGs from gut samples of farmed salmon were assembled in this study, including 2 MAGs from re-assembling previously published metagenomes^24^ and 68 MAGs from sequencing data generated in this study in 1) one sequencing run at NMBU and 2) three sequencing runs at SLU and SciLifeLab (Uppsala, Sweden). Libraries for the run at NMBU and the first run at SLU (August 2021) were prepared using the Nextera XT DNA Library Preparation kit (Illumina, San Diego, CA, USA) and paired-end 250 bp sequenced on an Illumina MiSeq. Libraries for the second (April 2022) and third run (January 2023) at SLU were prepared in-house using the Celero EZ DNA-Seq Modular Workflow v2 and Revelo DNA-Seq Enz for MagicPrep NGS v1 (Tecan, Männedorf, Switzerland), respectively, and paired-end 150 bp sequenced on an Illumina NovaSeq 6000 hosted by SciLifeLab. Raw reads were quality trimmed and adapter sequences removed using Trimmomatic v0.39^57^ (settings: PE -phred33 ILLUMINACLIP:adapters.fasta:2:30:10 LEADING:3 TRAILING:3 SLIDINGWINDOW:4:15 MINLEN:30).

To remove host contamination from both public and newly generated metagenome data, high-quality reads were mapped against the *Salmo salar* genome (GCF_000233375.1_ICSASG_v2) by minimap2^58^ using default parameters for short accurate genomic reads. Non-mapped paired reads were retrieved from minimap2 bam files using samtools v12^59^. Filtered data from each metagenome were assembled using MegaHit v 1.2.9 with the *“--no-mercy”* parameter. In addition, the filtered metagenomes (excluding those obtained by FACS) were co-assembled using the same parameters.

Metagenomic reads were mapped back to the respective metagenome assembly using minimap2 with the “-N 50 -ax sr” parameters. Samtools was used to produce sorted merged bam files. Bam files were then processed by the jgi_summarize_bam_contig_depths script from Metabat2 (“*-m 1500”* parameter) and binned with metaBat2 and MaxBin2 (“*-min_contig_length 900”* parameter) algorithms^60,61^. Completeness and contamination of each MAG was assessed with CheckM v1.2.0^62^ using the “lineage_wf” workflow.

### Taxonomy and functional annotation

All isolated assembled genomes and metagenome assembled genomes (MAGs) were classified taxonomically with GTDB-Tk-v-2.0.1^63^ (GTDB release 207). A maximum-likelihood tree was de-novo built using the protein sequence alignments generated by GTDB-Tk using IQ-Tree 2.0.3 with settings “*-m MFP -bb 1000 -nt 16”*, and the best amino-acid substitution model (LG+R5) was automatically selected by ModelFinder^64^ using the Bayesian Information criterion. A combined plot showing the phylogenetic tree, taxonomy and the isolation source (metagenomics or single culture), water source and the presence of 16S rRNA gene of all bacterial genomes in the atlas were produce using the GGTree Bioconductor package^65^ and an in-house made R script available at https://github.com/TheMEMOLab/MetaGVisualToolBox/blob/main/scripts/GenoTaxoTree.R

Functional annotation of the bacterial genomes, including gene prediction, ribosomal rRNA, tRNAs and gene annotation were performed by DRAM v 1.2.4^66^ with the following databases: Uniref90, PFAM-A, KOfam and dbCAN-V10 (all downloaded on Sep 2021). Antimicrobial resistance genes, toxins, virulence factors and mobile genetic elements were annotated using PathoFact v1.0^37^ with settings ‘workflow: “complete”, size_fasta: 10000, tox_threshold: 40, plasflow_threshold: 0.7, plasflow_minlen: 1000‘, and databases downloaded with the software in February 2023.

### Species-level clustering and pan-genome estimation

The 211 SMGA genomes were grouped into species-like clusters (mOTUs) based on a 95% ANI threshold using mOTUlizer^67^. Pan-genomes were computed for all mOTUs using mOTUpan from the same software package, whereby the MMseqs2^68^ amino acid identity threshold for clustering proteins was changed to 95%, and the coverage threshold was kept at the default 80%. The software uses a Bayesian method to infer core genomes (genes present in almost all genomes of the mOTU) and accessory genomes (genes present only in some genomes of the mOTU) that takes genome completeness estimates into account and is thus also applicable to incomplete genomes.

### Generation and analysis of strain-specific genome-scale metabolic reconstructions

The first step in creating genome-scale metabolic reconstructions for the sequenced microbes was to map the microbial genomes against the Kbase resource^69^. Kbase contains over 8,000 (draft) metabolic reconstructions, which included reconstructions for 94 of the 211 genomes. After downloading these draft reconstructions, we refined them using the DEMETER pipeline^70^. For three of these 94 microbes (*Photobacterium phosphoreum* sp. S39bc34, *Serratia liquefaciens* sp. S38bc38, and *Pseudomonas_E* sp. S3bc03), exo-metabolomic data was available, which was also used for the refinement with DEMETER^70^. Therefore, we identified those metabolites that could be consumed or secreted by the microbes by comparing untargeted metabolomics from chitin-supplemented minimal medium with and without a microbe. A metabolite was designated to be taken up if its concentration in the chitin-supplemented medium with the microbe was lower than its concentrations without the microbe and vice-versa.

After refinement of the microbial metabolic reconstructions, they were converted into condition-specific models by constraining to a nutrient unlimited and anoxic environment by setting the upper and lower flux bounds on the exchange reactions, which exchange metabolites to and from the extracellular environment, to 1000 and - 1000. The anoxic conditions were simulated by setting the lower flux bound of the oxygen exchange reaction, VMH ID: EX_o2(e), to zero. Next, the genome-scale metabolic models were interrogated using flux balance analysis (FBA)^71^. In FBA, an objective function, often biomass production, is either minimized or maximized, while assuming the system to be at a steady state, i.e., there is no change in concentration over time. The underlying linear optimization problem is formulated as follows:

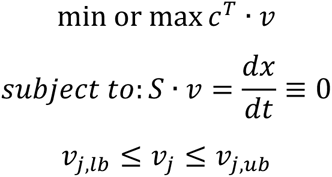

where c is a weight vector of zeros with one or more non-zero entries signifying the linear objective function. *v* is a flux vector to be solved for, containing a flux value for each on the n reactions in the model. *v_j,lb_* and *v_j,ub_* denominate the lower and upper flux bound for reaction *j* for all n reactions in the model. *S(m,n)* is the stoichiometric matrix, where the rows correspond to the mass-balances for each metabolite *i* and the columns correspond to the reactions. If a metabolite *i* participates in a reaction j the entry *S_i,j_* is non-zero and otherwise *S_i,j_* is zero.

We used parsimonious flux balance analysis (pFBA)^72^, a two-step FBA-based method, where first the objective function (here, biomass reaction; EX_bio1) is maximized. The resulting maximally possible flux value is then used in a second step to constrain the lower bound on the biomass reaction. Then, a quadratic optimization problem is solved, in which the total flux is minimised:

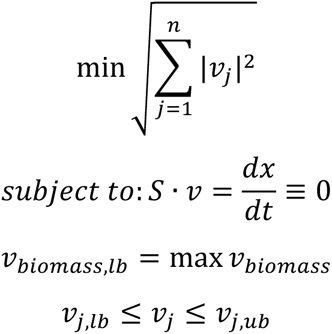

where

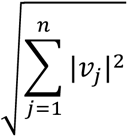

represents the Euclidean norm of the flux vector with size n for each *j* reaction. Minimization of the Euclidean norm results in a unique flux vector.

Additionally, we carried out flux variability analysis^73^ to compute the minimally and maximally possible flux value for each reaction in the model, while setting the lower bound of the biomass reaction to either 0 or 75% of the maximally possible flux value for the biomass reaction (**Supplementary Table S2)**. Again, an unlimited, rich, anoxic medium was simulated (see above). A metabolite could be secreted by the model if it carried a positive maximum flux and consumed if it carried a negative minimum flux value.

We then compared the predicted exo-metabolome, i.e., uptake and/or secretion of metabolites in the models with the measured exo-metabolomic data. First, we mapped the 121 measured metabolites onto the namespace of the metabolic models (i.e., onto the VMH database ^74^) using metabolite names and InchiKeys. In total, 57 metabolites could be mapped. Note that this does not mean that these 57 metabolites appear in all metabolic reconstructions. *In silico*, a metabolite can be taken up if the minimally possible flux value is negative or secreted if the minimally possible flux value is positive through the corresponding exchange reaction (denoted with an ‘EX_’, **Supplementary Table S2).**

All simulations were performed using the COBRA Toolbox^75^, with MATLAB 2020b as programming environment and IBM ILOG CPLEX 12.10 as linear and quadratic programming solver.

### SMGA’s 16S rRNA gene mapping to public amplicon datasets

16S amplicon datasets of Atlantic salmon gut samples were downloaded from NCBI (bioprojects: PRJEB39298, PRJNA498084, PRJNA555355, PRJNA590084, PRJNA594310, PRJNA650141, PRJNA730696, PRJNA733893, PRJNA824235, PRJNA824256, PRJNA866155) together with data derived from two in-house trials with salmons feeding on a commercial standard diet (PRJEB60544 and PRJEB60545). Fastq files in each Bioproject were downloaded using the fasterq-dump 3.0.0 tool from the SRA toolkit^76^ and Fastp was used to inspect reads for sequencing adapters and perform quality trimming (q >25). The DADA2 pipeline^77^ was then used for reads denoising, merging and screening for chimeric sequences, which were subsequently removed, to finally produce amplicon sequence variants (ASVs) of each Bioproject. ASVs were compared to a database of SMGA 16S rRNA gene sequences, complete as well as partial sequences covering the amplified regions in the used amplicon datasets, which were present in 146 of the 211 SMGA genomes. We obtained 531 distinct sequences from these 146 genomes. Comparison to all ASVs obtained from the amplicon datasets was done using ncbi-blast-2.13.0+^78^ with settings ‘-task “blastn”’ and ‘-max_target_seqs 531’ (number of sequences in the database). Hits were filtered for ≥97% identity (“pident”) and ≥95% query coverage (“qcovhsp”). For each SMGA genome, the best of the remaining hits (if any) to each amplicon dataset was extracted. Results were plotted using ggplot2 v3.4.0^79^ in R v4.2.2^80^.

### Metatranscriptomics analysis using the SMGA as a database

Midgut content samples were obtained following dissection of 33 salmons that were fed with a standard commercial diet and raised at 12 °C in freshwater (T0, T1 and T2) and transferred to seawater (T3) at EWOS Innovation Center, Dirdal, Norway. All samples were preserved in DNA/RNA SHIELD™, obtained by Zymo Research, following the Zymo Research standard procedure. RNA was extracted using the methods reported in^81^. RNA concentration and purity were determined using a Qubit 3.0 fluorometer and a Nanodrop 8000 (Thermo Scientific, Wilmington, USA). RNA integrity was checked by using an Agilent 2100 Bioanalyzer (Agilent Technologies, Santa Clara, CA, USA). Prior to analysis, all samples were randomized. Library preparations were carried out by Novogene (Beijing, China) using a TruSeq Stranded mRNA kit (Illumina, San Diego, CA, USA), as per manufacturer’s protocol. Libraries were sequenced on the lllumina NovaSeq 6000 platform at Novogene, (Beijing, China), using 300 bp paired-end sequencing. Three extraction negatives and two library negatives were included. The resulting sequence reads were filtered for quality using fastp v. 0.12.4 with an average Phred threshold of 30 (-q 30). rRNAs and tRNAs was removed from the reads using SortMeRNA v 4.3.4^82^ with the following Silva databases: silva-bac-16s-id90, silva-arc-16s-id95, silva-bac-23s-id98, silva-arc-23s-id98, silva-euk-18s-id95, silva-euk-28s-id98 and the parameters: --out2 --paired_out –fastx --thread 12. To remove all sequences derived from the host, the filtered reads were aligned to the *Salmo salar* genome Ssal_v3.1 RefSeq ID GCF_905237065.1 using the STAR v 2.7.9a alignment suit^83^. All non-mapped reads were retrieved from the sam files using Samtools 1.13 and the parameters -f 12 -F 256 -c 7. These reads were used to quantify the expression of ORFs encoded by the SMGA genomes and MAGs using kallisto v 0.44.0. The resulting transcripts per million (TPM) abundance tables of each metatranscriptomic sample were gathered into a single table using the Bioconductor tximport 1.26.1 library in R 4.2.3. Principal component analysis (PCA) was implemented to visualise samples clustering and manual removing of outliers resulting in 33 samples for further detection of bacterial gene expression. A bacterial gene from the SMGA was considered expressed if it shows a value higher than one TMP in at least one replicate of the experiment. Variation in the SMGA bacterial gene expression among samples were visualized in terms of Z-scores in a heatmap generated using the pheatmap function in R. The data was used to reconstruct the active metabolic pathways displayed in **Fig. 4a**.

### Untargeted metabolomic analysis

Triplicate cultures of Photobacterium *phosphoreum* sp. S39bc34, Serratia *liquefaciens* sp. S38bc38 and *Pseudomonas_E* sp. S3bc03 were grown at 18 °C overnight in M9 medium ((8 g/L Na_2_HPO_4_, 4 g/L KH_2_PO_4_, 0.5 g/L NaCl, 0.5 g/L NH_4_Cl, 0.5 g/L EDTA, 0.083 mg/L FeCl_3_ x 6 H_2_O, 0.863 mg/L ZnCl_2_, 0.013 mg/L CuCl_2_ x 2 H_2_O, 0.1 mg/L CoCl_2_ x 6 H_2_O, 0.1 mg/L H_3_BO_3_, 0.016 mg/L MnCl_2_ x 6 H_2_O, 1 mM MgSO_4_, 0.3 mM CaCl_2_, 1 mM thiamine hydrochloride, 1 mM biotin, 0.5% [wt/vol] beef extract from Sigma-Aldrich, St. Louis, MO, USA) supplemented with 5% (wt/vol) glucose (Sigma-Aldrich). Overnight cultures were then used to inoculate M9 medium supplemented with 5% (wt/vol) chitin from shrimp shells (Sigma-Aldrich) and grown at 18 °C for 48-72 h. Uninoculated M9 media with chitin were also incubated as a negative control group and this group had three biological replicates. Cells were harvested by centrifugation at 16,000 x g for 5 minutes and culture supernatant processed for semi-polar metabolite analysis. Sample analysis was carried out by MS-Omics (Vedbæk, Denmark) using a UPLC system (Vanquish, Thermo Fisher Scientific) coupled with a high-resolution quadrupole-orbitrap mass spectrometer (Orbitrap Exploris 240, Thermo Fisher Scientific). The UHPLC was performed using an ACQUITY UPLC HSS T3 C18 lined column, with dimensions of 2.1 × 150 mm and a particle size of 1.8 µm. The composition of mobile phase A was 10 mM ammonium formate at pH 3.1 in 0.1% formic acid LC–MS grade (VWR Chemicals) and 10 % ultra-pure water (Merck KGaA). The mobile phase B contained 10 mM ammonium formate at pH 3.1 in 0.1% formic acid in methanol. The flow rate was kept at 300 µl ml−1 consisting of a 2 min hold at 0% B, increased to 35% B at 12 min, increased to 90% B at 13 min and held for 1 min, and finally decreased to 0% B at 15 min. The column temperature was set at 30 °C and an injection volume of 50 µl was used. Surfactant removed samples (using zinc nitrate hexahydrate and ammonium thiocyanate) were diluted 10 times in mobile phase eluent A and fortified with stable isotope labelled standards before analysis before injection. An electrospray ionization interface was used as an ionization source. Analysis was performed in positive and negative ionization mode under polarity switching. Data were processed using Compound Discoverer 3.3 (ThermoFisher Scientific) and Skyline 21.2. Identification of compounds were performed at four levels; Level 1: identification by retention times (compared against in-house authentic standards), accurate mass (with an accepted deviation of 3 ppm), and MS/MS spectra, Level 2a: identification by retention times (compared against in-house authentic standards), accurate mass (with an accepted deviation of 3 ppm). Level 2b: identification by accurate mass (with an accepted deviation of 3 ppm), and MS/MS spectra, Level 3: identification by accurate mass alone (with an accepted deviation of 3 ppm).

## Data availability

Oxford Nanopore sequencing reads have been deposited in the sequence read archive SRA with project numbers PRJEB45024 and PRJEB61648. Amplicon sequencing reads are available under SRA BioProjects PRJEB60544 (ImpTrial2) and PRJEB60545 (ImpTrial1). Shotgun metagenomic reads have been deposited at SRA BioProjects PRJEB60591 and PRJNA947914. RNA sequencing reads can be found in SRA under accession number PRJEB60552. Mass spectrometry data for this study can be found on the Mass Spectrometry Interactive Virtual Environment (MassIVE) repository (massive.ucsd.edu) with accession number MSV000089895. The SMGA is publicly available via Figshare (Genomes_fasta, 10.6084/m9.figshare.22691881; Genes_nuc_fna, 10.6084/m9.figshare.22691869; Genes_prot_faa, 10.6084/m9.figshare.22691980).

## Acknowledgements

This work was supported by the Research Council of Norway (project no. 300846), the Swedish Research Council Formas (grant no. 2019-02336) and the European Union’s Horizon 2020 research and innovation program under the ERA-Net Cofund project BlueBio (grant agreement no. 311913). IT received funding from the European Research Council (ERC) under the European Union’s Horizon 2020 research and innovation programme (grant agreement no. 757922) and from the Science Foundation Ireland under Grant number 12/RC/2273-P2. Sequencing was performed by the SNP&SEQ Technology Platform in Uppsala, part of the National Genomics Infrastructure (NGI) Sweden and SciLifeLab. Cell sorting and whole genome amplification was performed at the Microbial Single Cell Genomics Facility (MSCG) at SciLifeLab. Computations were performed on resources provided by the Swedish National Infrastructure for Computing (SNIC) at Uppsala Multidisciplinary Center for Advanced Computational Science (UPPMAX) under projects SNIC 2021/5-51 and SNIC 2021/22-602. The Orion High Performance Computing Center at the Norwegian University of Life Sciences and Sigma2 - the National Infrastructure for High Performance Computing and Data Storage in Norway are acknowledged for providing computational resources that have contributed to meta-omics analyses described in this study. We acknowledge Claudia Bergin at the SciLifeLab Microbial Single Cell Genomics Facility for support with cell sorting, genome amplification and library preparation.

## Author contributions

P.B.P., S.L.L.R., A.V.P.L, S.R.S., S.B. and T.R.H. designed the study. Isolation experiments and genomics analysis of the cultured microbes were carried out by A.V.P.L., S.L.L.R. and S.M.J. Metagenomic analyses were performed by A.V.P.L. and M.H. Culture experiments and untargeted metabolomic analyses were carried out by S.L.L.R. Constraint-based metabolic models were generated by T.H., B. W. and I.T. M.H. and S.G. conducted amplicon sequencing analyses. C.R.K., K.R., L.S., J.A.R., M.T.L. and S.B. obtained isolates and generated shotgun sequencing data. The draft manuscript was written by S.L.L.R., P.B.P., M.H., A.V.P.L. and T.H. All authors contributed to the editing of the text and content and approved the final version.

## Supplementary Material

**Supplementary Fig. S1.**
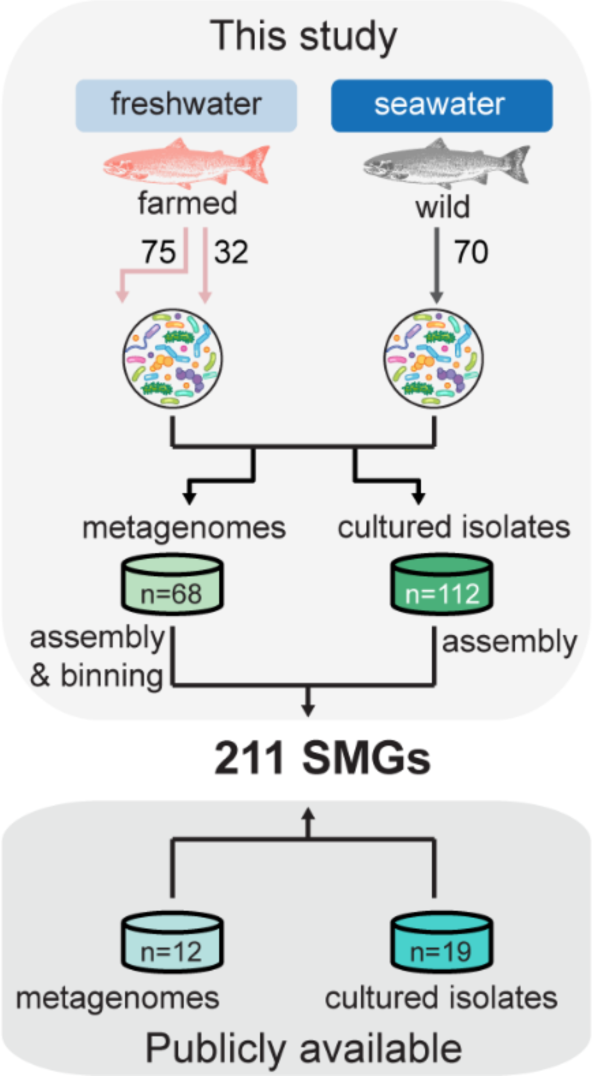
Strategy for the generation of the SMGA. Digesta samples were collected from 107 farmed and 70 wild fish either at the freshwater or seawater stage. Genomic and metagenomic datasets were combined to generate a collection of 211 salmon gut microbial genomes. Green boxes indicate the number of genomes from bacterial isolates or bacterial MAGs obtained using two different approaches in this study. Turquoise boxes indicate the number of genomes for cultured isolates or bacterial MAGs from publicly available studies. For a description of the different assembly strategies, see the Methods section.

**Supplementary Fig. S2.**
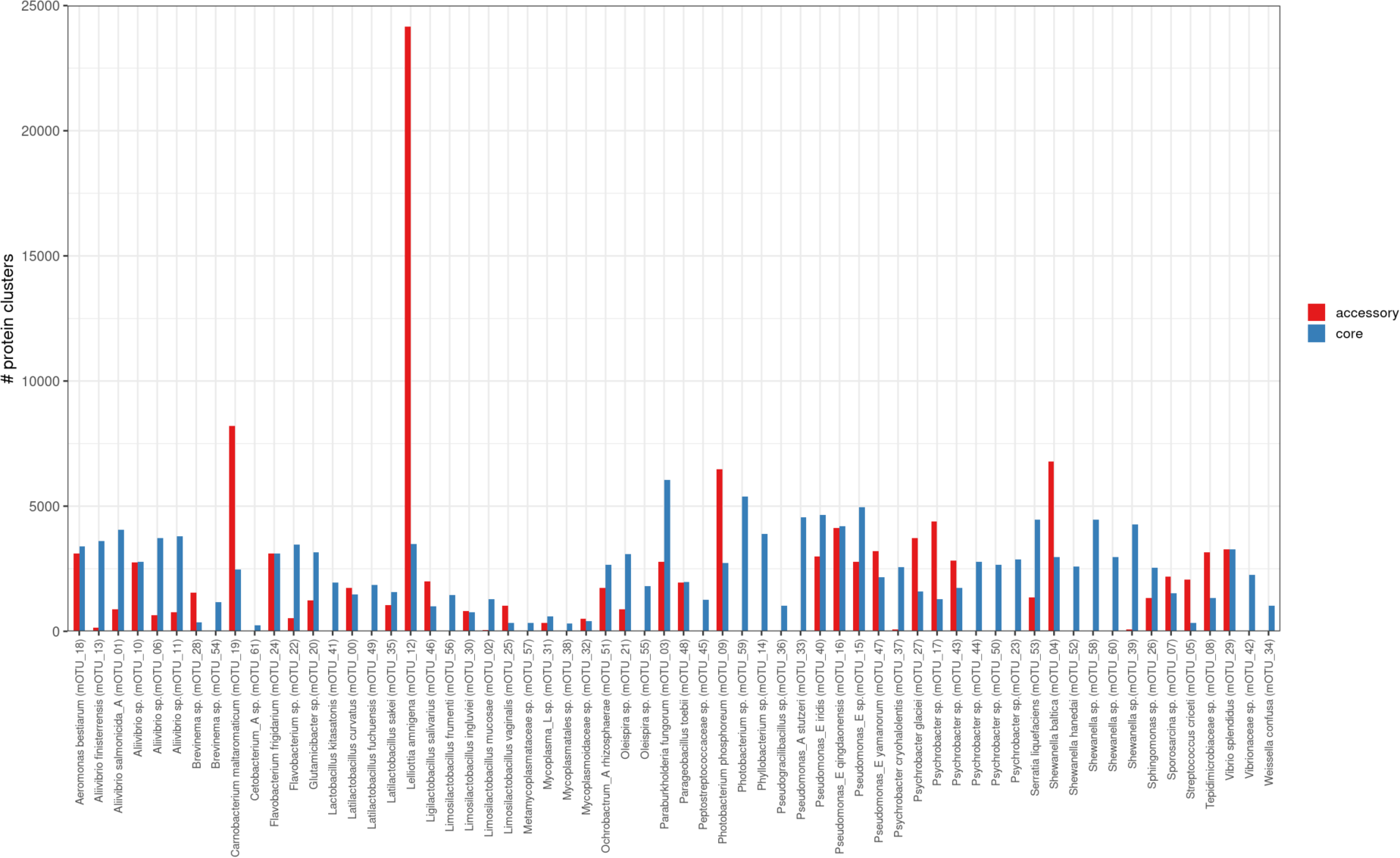
Pan- genome sizes of species-like mOTUs. The 211 SMGA genomes were clustered into 62 mOTUs (x-axis) based on 95% ANI. Bar heights indicate the number of protein clusters within the core and accessory genome of each mOTU.

**Supplementary Fig. S3.**
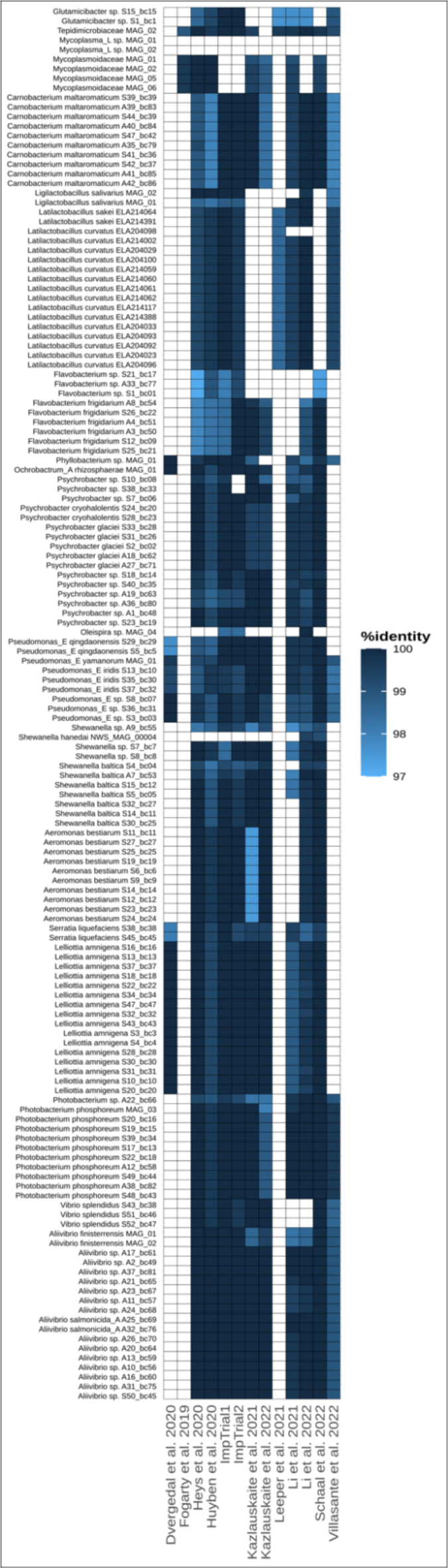
Detecting genomes from the SMGA in publicly available datasets. The detection of isolate genomes and MAGs from the SMGA (y-axis) in selected publicly available 16S rRNA gene amplicon datasets (x-axis) based on alignment of 16S rRNA gene sequences (y-axis). 16S rRNA gene detection is coloured based on the % identity of the gene alignment. At a 97% identity level to amplicon sequence variants (ASVs), 144 out of 146 SMGA bacteria were detected in publicly available 16S rRNA gene datasets from either *in vivo* trials or *in vitro* models with salmon gut microbial communities.

**Supplementary Fig. S4.**
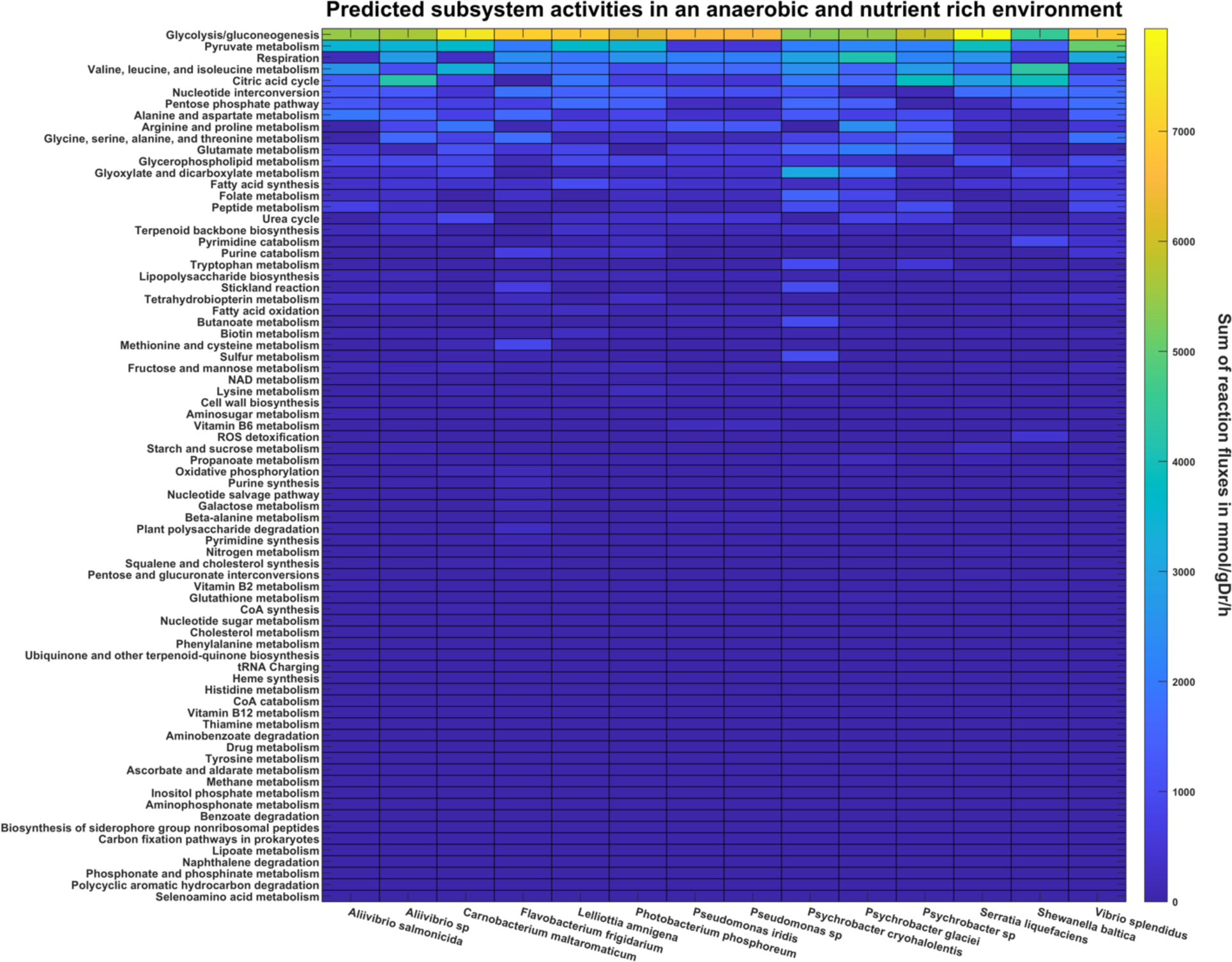
Heatmap of predicted total metabolic flux in each metabolic subsystem. The total metabolic flux for each subsystem was calculated by taking the absolute sum of all flux values of reactions within a subsystem. The species-specific reaction fluxes represent the mean average metabolic flux of all created strain models within a microbial species. All models were interrogated in an anaerobic and nutrient rich environment, meaning that all nutrients needed for growth were available in sufficient quantities for the models to produce biomass. All species predicted largely similar subsystem activities. The most active subsystems include those involved in energy metabolism, nucleotide metabolism, and amino acid metabolism.

**Supplementary Fig. S5.**
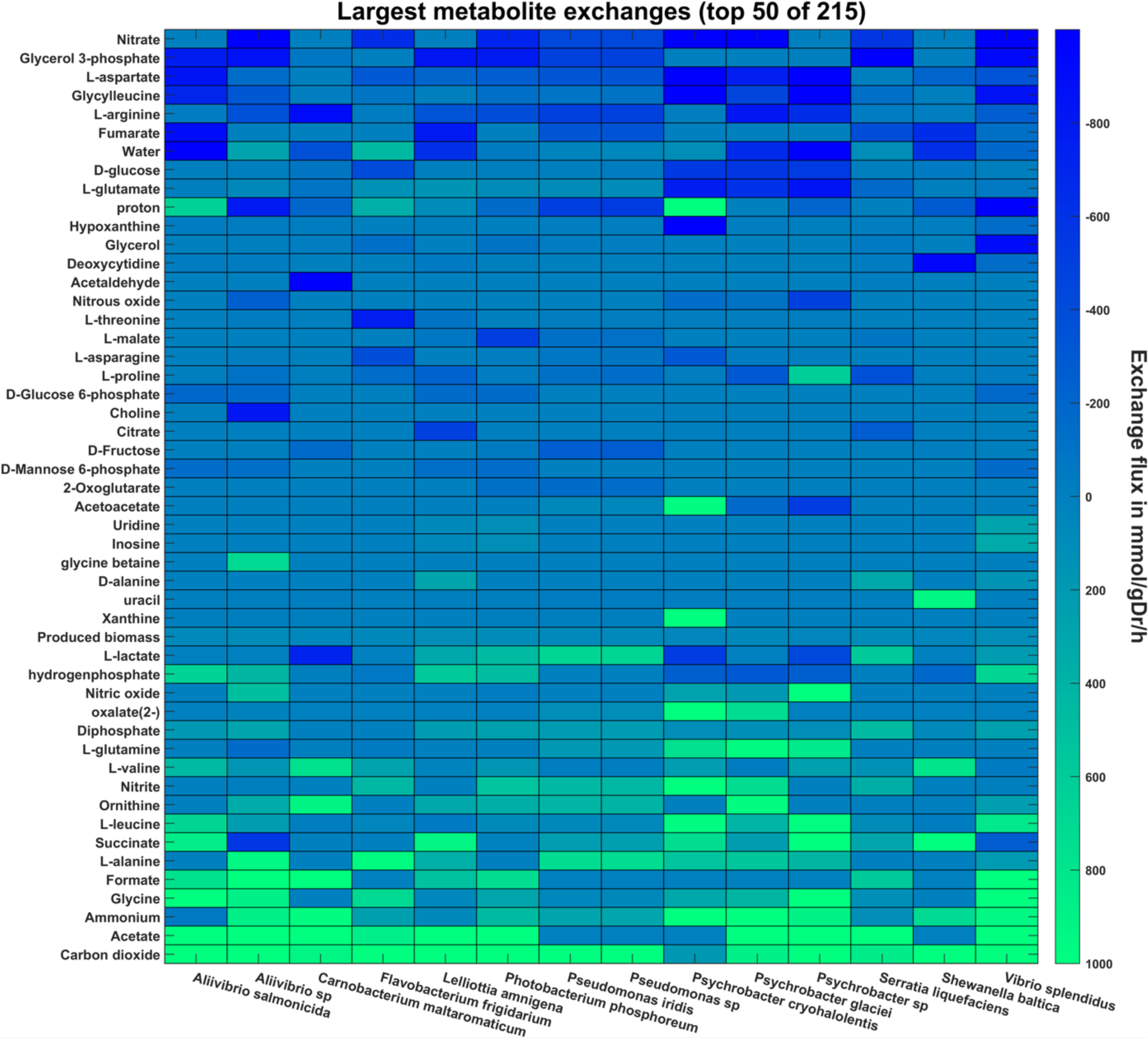
Heatmap of the largest 50 predicted metabolite exchanges for each species. The species-specific reaction fluxes represent the mean average metabolic flux of all created strain models within a microbial species. An exchange flux below zero indicates that the metabolite is taken up by the system, while an exchange flux above zero indicates an excretion of the metabolite. Metabolites that are taken up in larger quantities by all models include nitrate, glycerol-3-phosphate, fumarate, and glucose. Carbon dioxide, acetate, and ammonium on the other hand, are among the most excreted metabolites.

**Supplementary Fig. S6.**
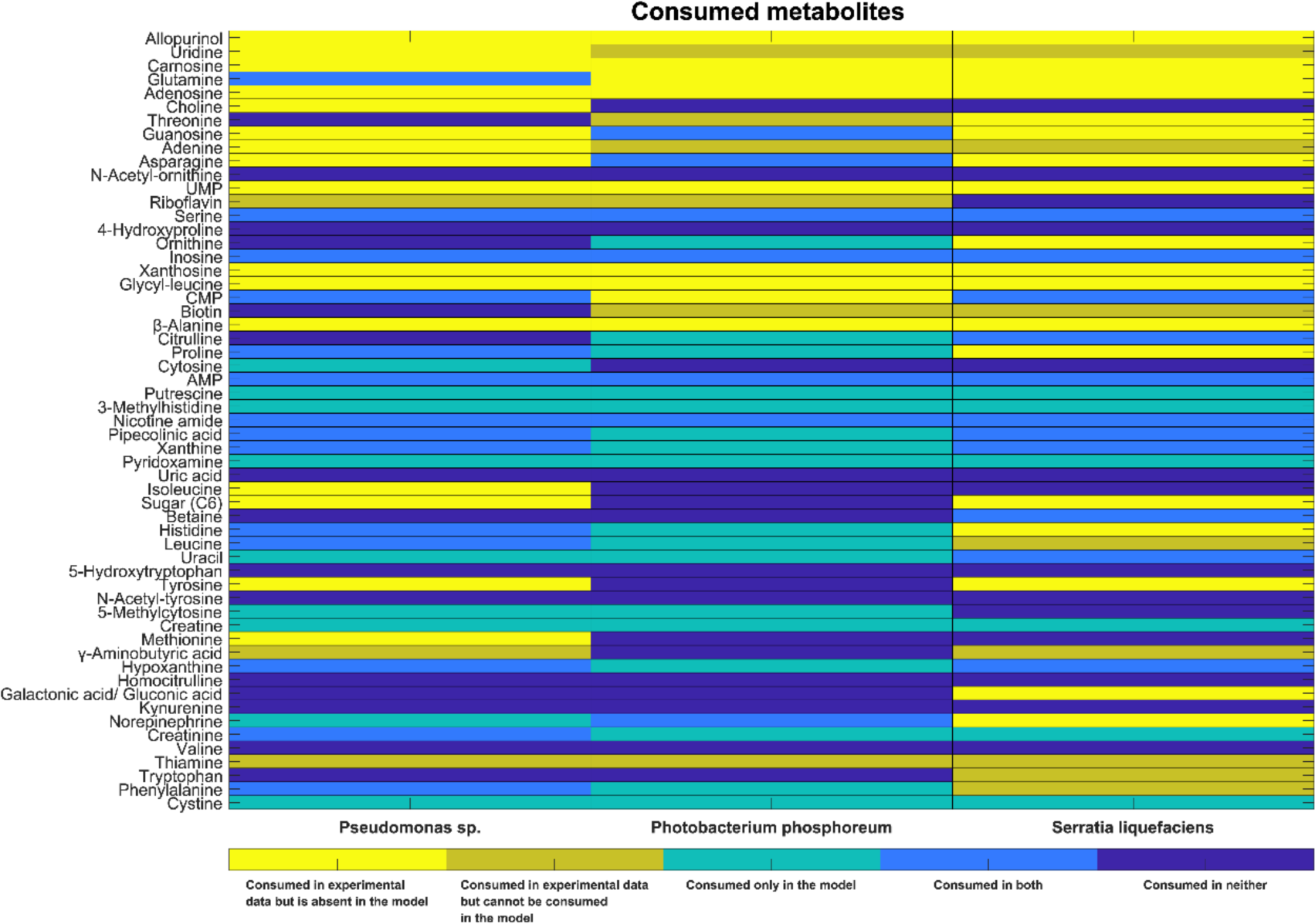
Heatmap summarising the metabolomic results and the genome-scale metabolic model capabilities. Only metabolites that could be mapped onto a VMH ID are shown. The yellow cells indicate that a metabolite was consumed in the experimental data, but was not present in the corresponding model. Gold cells indicate that a metabolite was consumed in the experimental data and was present in the model, but could not be consumed by the model. Turquoise cells indicate metabolites that were consumed by the model, but not in the experimental data, whereas blue cells indicate metabolites that were consumed in both the model and the experimental data. Dark blue cells indicate metabolites that could not be consumed in either the experimental data or the models.

**Supplementary Fig. S7.**
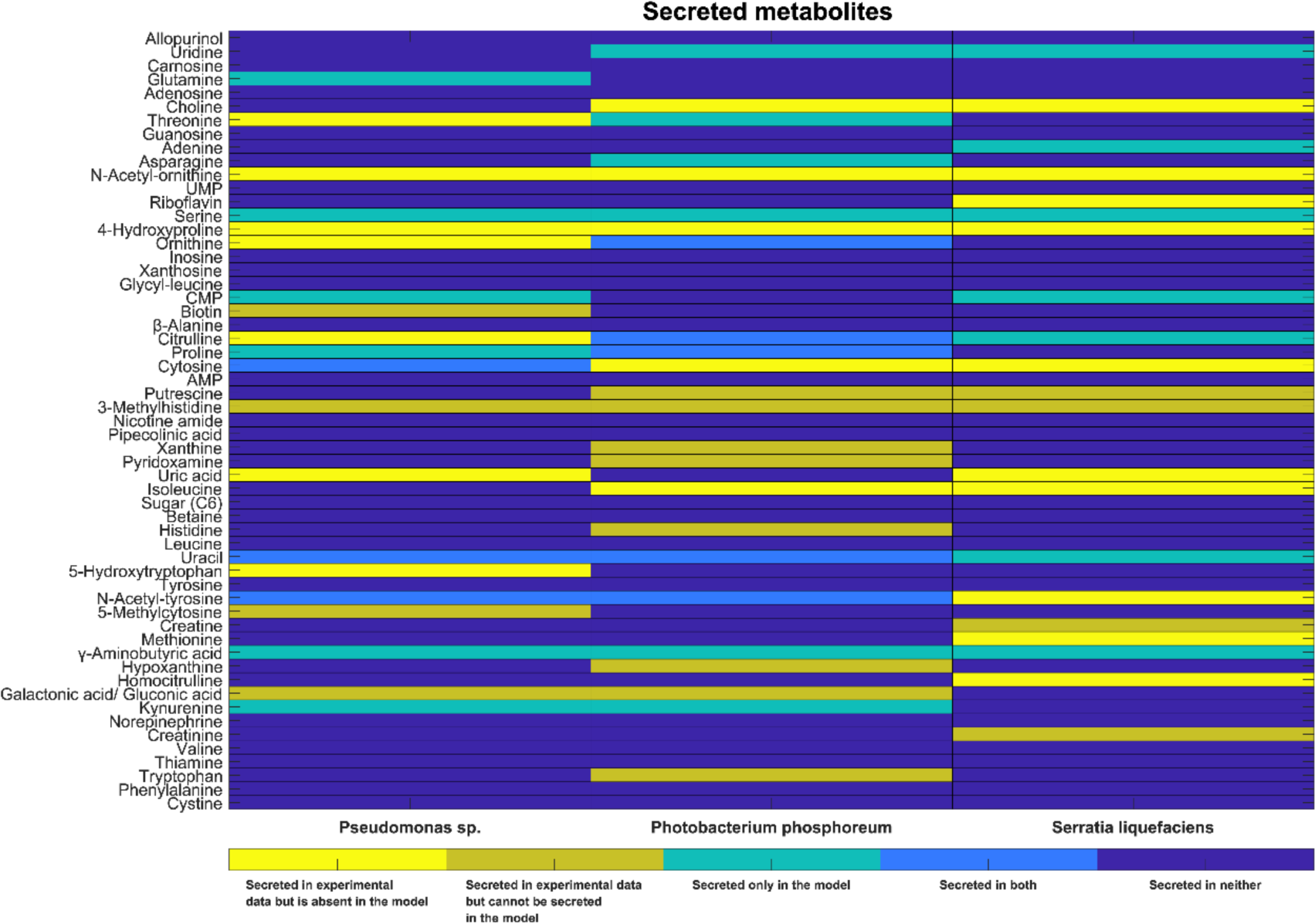
Heatmap summarising the exo-metabolomic results and the genome-scale metabolic model capabilities. Only metabolites that could be mapped onto a VMH ID are shown. The yellow cells indicate that a metabolite was secret in the experimental data but was not present in the corresponding model. Gold cells indicate that a metabolite was secreted in the experimental data and was present in the model but could not be secreted by the model. Turquoise cells indicate metabolites that were secreted by the model, but not in the experimental data, whereas blue cells indicate metabolites that were secreted in both the model and the experimental data. Dark blue cells indicate metabolites that could not be secreted in either the experimental data or the models.

**Supplementary Fig. S8.**
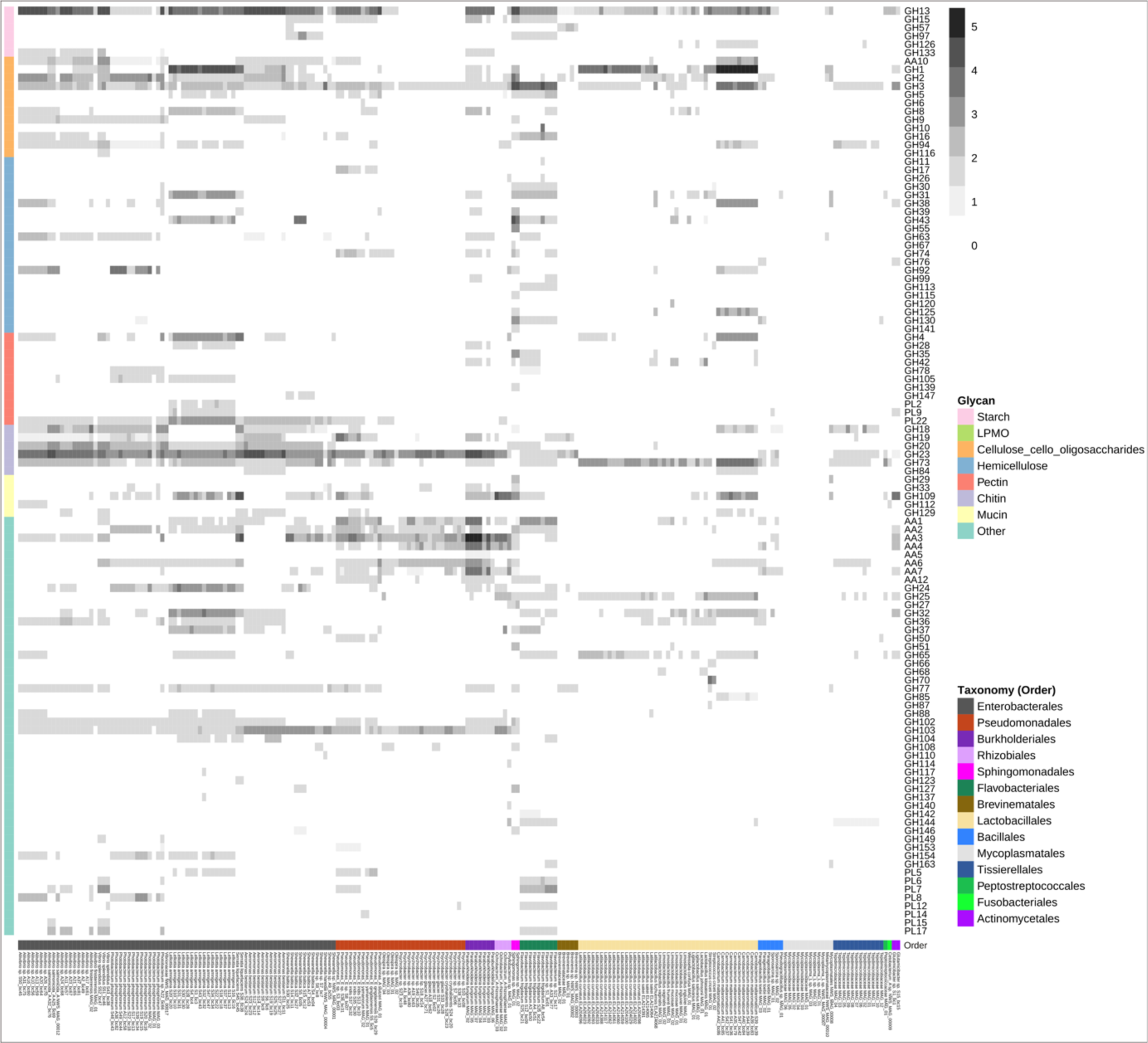
CAZyme profiles of the 211 salmon microbial genomes and MAGs in the SMGA. Heatmap showing the presence of CAZy families (listed on the righthand y-axis) arranged into the different glycan substrate categories (listed on lefthand y-axis) found in each genome that are arranged in taxonomic orders (x-axis). The presence of CAZy genes is denoted by grey-black boxes that are weighted for copy number. CAZy families that are not detected are represented by a white box. GH: glycoside hydrolase, PL: polysaccharide lyase, AA: auxiliary activity.

**Supplementary Fig. S9.**
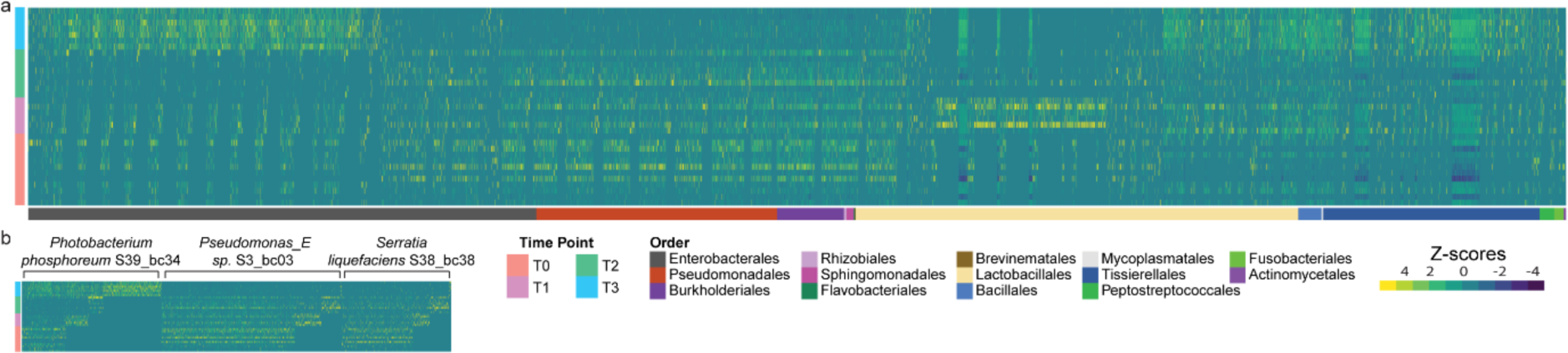
Heatmap illustrating the application of the SMGA as a database to map metatranscriptomes from gut samples. a) Variation in gene expression of all bacterial genes in the SMGA database (x-axis) and b) a subset of three Enterobacterales (*Photobacterium phosphoreum* S39_bc34, *Pseudomonas_*E sp. S3_bc03 and *Serratia liquefaciens* S38_bc38) in metatranscriptomes generated from gut samples obtained from 33 growing fish fed a standard commercial diet and collected at different life stages (y-axis). T0: 30 g fish (parr), freshwater; T1, 90 g fish (pre-smolt), freshwater; T2, 130 g fish (smolt), freshwater; T3, 300 g fish (adult), seawater.

**Supplementary Fig. S10.**
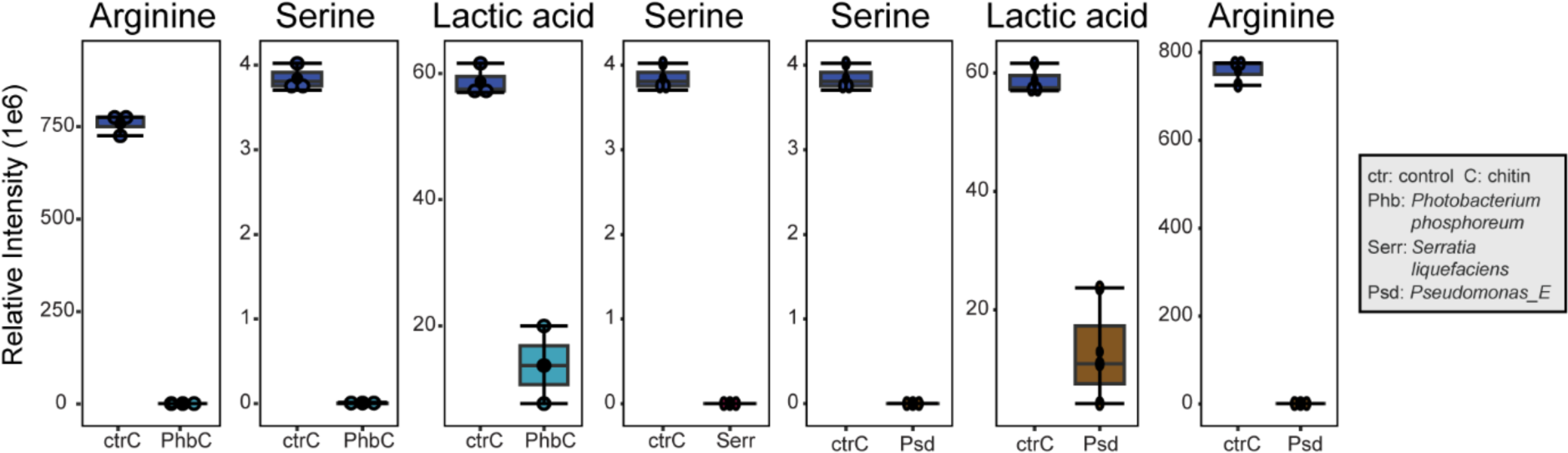
Untargeted metabolomics results for serine, arginine and lactic acid amounts in the spent supernatant of triplicate cultures growing on chitin.

